# Divergent encoding of active avoidance behavior in corticostriatal and corticolimbic projections

**DOI:** 10.1101/2021.09.15.460552

**Authors:** Bridget L. Kajs, Adrienne C. Loewke, Jeffrey M. Dorsch, Leah T. Vinson, Lisa A. Gunaydin

**Author notes:** These authors contributed equally (Bridget L. Kajs, Adrienne C. Loewke).

## Abstract

Active avoidance behavior, in which an animal performs an action to avoid a stressor, is crucial for survival and may provide insight into avoidance behaviors seen in anxiety disorders. Active avoidance requires the dorsomedial prefrontal cortex (dmPFC), which is thought to regulate avoidance via downstream projections to the striatum and amygdala. However, the endogenous activity of projection-defined dmPFC subpopulations during active avoidance learning remains unexplored. Here we utilized fiber photometry to record from the dmPFC and its downstream projections to the dorsomedial striatum (DMS) and the basolateral amygdala (BLA) during active avoidance learning in mice. We examined neural activity during conditioned stimulus (CS) presentations, active avoidance, and cued freezing. Both prefrontal projections showed learning-related increases in activity during CS onset throughout active avoidance training. The dmPFC as a whole showed increased activity during avoidance and decreased activity during cued freezing. Finally, dmPFC-DMS and dmPFC-BLA projections showed divergent encoding of active avoidance behavior, with the dmPFC-DMS projection showing increased activity and the dmPFC-BLA showing decreased activity during active avoidance. Our results identify differential prefrontal encoding of active and passive coping behaviors in the same behavioral paradigm and demonstrate divergent encoding of active avoidance in projection-specific dmPFC subpopulations.

## INTRODUCTION

Active avoidance is a behavioral coping strategy in which an organism performs an action to avoid a stressor and can be adaptively enacted to evade danger and ensure survival. However, active avoidance can become maladaptive when used in excess or in response to overexaggerated perceived threats as seen in anxiety disorders. Despite its high clinical relevance, our understanding of the neurobiological basis of active avoidance has lagged far behind other behaviors relevant to anxiety disorders such as approach-avoidance decision making or fear learning (LeDoux et al., 2017). The dorsomedial prefrontal cortex (dmPFC) is an attractive candidate to explore in the context of active avoidance given its clear ties to anxiety disorder pathophysiology (Holzschneider & Mulert, 2011; Rauch & Shin, 2002) and avoidance behavior in humans (Collins et al., 2014; Delgado et al., 2009) as well as clinically relevant behaviors in rodents (Giustino & Maren, 2015; Tovote et al., 2015). In non-psychiatric populations, dmPFC activity is associated with active avoidance learning (Collins et al., 2014) while in post-traumatic stress disorder (PTSD) patients, dmPFC activation during fear extinction positively correlates with patients’ avoidance symptoms (Sripada et al., 2013). In rodents, dmPFC plays a crucial role in associative fear learning (Adhikari et al., 2015; Corcoran & Quirk, 2007; Courtin et al., 2014; Dejean et al., 2016; Fenton et al., 2014; Giustino & Maren, 2015; Herry & Johansen, 2014; Klavir et al., 2017; Marek et al., 2018; Meyer et al., 2019; Sharpe & Killcross, 2014; Sierra-Mercado et al., 2011; Sotres-Bayon et al., 2012; Tovote et al., 2015) and instrumental action-outcome learning (Gourley & Taylor, 2016; Grace et al., 2007; Peters et al., 2005; Pinto & Dan, 2015), both of which are components of active avoidance behavior. Studies have directly demonstrated the importance of the dmPFC for a variety of avoidance behaviors including real time and conditioned place avoidance (Huang et al., 2020; Lee et al., 2014; Vander Weele et al., 2018), inhibitory avoidance (Garrido et al., 2012; Ito & Morozov, 2019; Izquierdo et al., 2007; Torres-García et al., 2017; Zhang et al., 2011), approach-avoidance decision making (Friedman et al., 2015; Loewke et al., 2021), and active avoidance (Beck et al., 2014; Bravo-Rivera et al., 2014; Capuzzo & Floresco, 2020; Diehl et al., 2018, 2020). One recent study using the platform-mediated active avoidance task showed that suppression of dmPFC activity is associated with avoidance learning (Diehl et al., 2018). Another study using a discriminative two-way active avoidance paradigm found that dmPFC population activity alone could be used to decode conditioned stimulus (CS) identity between a conditioned stimulus that predicted shock and led to robust avoidance behavior (CS+) and a conditioned stimulus that did not predict to shock and did not lead to avoidance (CS-) (Jercog et al., 2021). In these studies, task-relevant neural activity in the dmPFC during active avoidance has only been examined on the final day of active avoidance training after learning has already occurred. To our knowledge, no studies have thoroughly examined dmPFC activity throughout avoidance learning. Investigating how task-relevant signals in the dmPFC develop in real time across days of learning could help determine whether the dmPFC is preferentially recruited during certain stages of learning or whether task-relevant dmPFC activity requires consolidation across days.

Further dissecting the dmPFC into subpopulations based on their projection target may also yield more refined insights into the nuanced and varied roles of the dmPFC in active avoidance behavior. One downstream target of the dmPFC that has been consistently tied to active avoidance behavior is the basolateral amygdala (BLA) (Amorapanth et al., 2000; Bravo-Rivera et al., 2014; Choi et al., 2010; Darvas et al., 2011; Diehl et al., 2020; Killcross et al., 1997; Kyriazi et al., 2018; Lázaro-Muñoz et al., 2010; Maren et al., 1991; Poremba & Gabriel, 1999). The BLA has subpopulations of cells that specifically encode successful active avoidance behavior (Kyriazi et al., 2018), and inactivating the BLA impairs platform-mediated active avoidance behavior (Bravo-Rivera et al., 2014). Additionally, the dmPFC-BLA projection has been directly tied to active avoidance, as optogenetically stimulating or inhibiting this projection bidirectionally affects platform-mediated active avoidance behavior (Diehl et al., 2020). However, despite these optogenetic studies suggesting a causal role of this projection in avoidance behavior, no studies have directly recorded the endogenous activity in this projection subpopulation during active avoidance learning or expression. The task-relevant information of the dmPFC-BLA projection in active avoidance could be multifold. dmPFC-BLA projections could signal crucial information about the cue-shock association, as the BLA receives associative information that has converged upstream in the lateral amygdala (LA) (Duvarci & Pare, 2014; Tovote et al., 2015). dmPFC-BLA projections may also directly impact behavioral output by amplifying avoidance information sent to the nucleus accumbens (LeDoux et al., 2017; Ramirez et al., 2015) and suppressing fearful freezing information sent to the central amygdala (LeDoux et al., 2017; Terburg et al., 2018). However, it remains unknown how the real-time neural dynamics in this projection encode active avoidance, which would require projection-specific recording of dmPFC-BLA projection neurons during avoidance learning and expression.

While corticolimbic projections have been heavily studied in the context of fear conditioning, recent evidence suggests that corticostriatal projections also play a key role in avoidance behavior (Friedman et al., 2015; Loewke et al., 2021). Human fMRI studies have implicated both the dorsal and ventral striatum in active avoidance behavior (Boeke et al., 2017; Collins et al., 2014; Delgado et al., 2009). However, while the ventral striatum has been more thoroughly studied in rodent models (Bravo-Rivera et al., 2014, 2015; Darvas et al., 2011; Gentry et al., 2016; Oleson et al., 2012; Piantadosi et al., 2018; Ramirez et al., 2015; Rodriguez-Romaguera et al., 2016; Stelly et al., 2019; Wenzel et al., 2018), there has been less exploration into the role of the dorsal striatum in active avoidance (Boschen et al., 2011; Dombrowski et al., 2013; Wendler et al., 2014; Wietzikoski et al., 2012). dmPFC projections to the dorsal striatum, especially the dorsomedial subregion (DMS), are uniquely positioned to play a crucial role in active avoidance behavior given their importance in goal-directed behavior (Balleine & O’Doherty, 2010; Gremel & Costa, 2013; Hart, Bradfield, & Balleine, 2018; Hart, Bradfield, Fok, et al., 2018; Pitts et al., 2018) and approach-avoidance decision making (Friedman et al., 2015; Loewke et al., 2021). Additionally in humans, the degree of coupling between the caudate (the human homologue of the DMS) and the medial prefrontal cortex (mPFC) positively correlates with successful active avoidance performance with greater coupling predicting better performance (Collins et al., 2014). The dmPFC-DMS projection could hold task-relevant information regarding action-outcome contingencies necessary for goal-directed behavior (Balleine & O’Doherty, 2010; Yin & Knowlton, 2006). As the dmPFC-DMS projection directly interfaces with the striatum, this projection could also carry crucial information for avoidance initiation through movement-promoting pathways (Kravitz & Kreitzer, 2012; Redgrave et al., 2010). Despite promising initial evidence and strong rationale for its involvement, the dmPFC-DMS projection has remained completely unexplored in rodent models of active avoidance.

In this study, we utilize fiber photometry in combination with retrograde viral targeting strategies to examine the activity of the dmPFC and its projections to the DMS and the BLA during learning and expression in a cued active avoidance task. We identified task-relevant neural activity in response to CS onset as well as clinically relevant behaviors such as avoidance and freezing. We find that dmPFC and both of these downstream projections show learning-related increases in activity at CS onset. However, encoding by these projections diverges during avoidance onset, where we find increased activity in the dmPFC-DMS projection and decreased activity in the dmPFC-BLA projection. Finally, we identify decreases in dmPFC activity that correspond to freezing bouts. Overall, our results suggest that dmPFC and its projections to DMS and BLA contain task-relevant information and that the dmPFC-DMS and dmPFC-BLA may play distinct yet complementary roles in successful enactment of active avoidance behavior.

## RESULTS

### dmPFC shows learning related increases in activity at CS onset

To record the endogenous activity of excitatory dmPFC neurons during avoidance learning, we utilized a virally-expressed calcium indicator (GCaMP) and fiber photometry to record changes in GCaMP fluorescence in the dmPFC, which acted as a proxy for changes in neural activity (**Figure 1A, Supplemental Figure 1**). Mice were trained for five days on a cued two-way active avoidance behavioral paradigm (**Figure 1B**). A white light underneath the shock floor where the animal was present acted as a conditioned stimulus (CS) and signaled impending shock on that side of the two-chamber apparatus. Throughout training, animals learned to successfully avoid the impending shock by shuttling from the lit chamber to the unlit chamber during the CS-only period. Animals were trained until they successfully avoided the shock on 80% of all trials, which occurred by day 5 of training (**Figure 1C**). Average avoidance latency was between 4-6 seconds and decreased across training. Avoidance latencies also became more stereotyped as evidenced by a change in the shape of the avoidance latency distribution from a broad non-specific curve on day 1 to a narrower distribution on day 5 (**Figure 1D**). To uncover task-relevant neural activity in the dmPFC during active avoidance learning, we first examined heatmaps of the average change in calcium signal in the dmPFC for each trial during the CS-only period (first 10 seconds of the CS before the shock occurred) (**Figure 1E**). We saw a rapid peak in fluorescence at CS onset on day 1 that occurred on most but not all trials and became more consistent throughout training. In addition to this rapid CS response, we also observed a sustained increase in fluorescence across the 10 second CS-only period that appeared to develop across learning, as it was consistently present on days 3 and 5 but not day 1. While these heatmaps provided initial insight that there were task-relevant changes in the dmPFC during active avoidance learning, it was unclear whether these changes in calcium signal were a response to the CS itself and/or represented behaviors such as avoidance or freezing. In order to isolate CS onset responses, we created a perievent time histogram (PETH) of z-scored changes in dmPFC calcium signal during the first second of the CS presentation as the majority of avoidance movements (>90%) occurred after this time window (**Figure 1F**). We found that the dmPFC showed a sharp increase in fluorescence during the first second of CS onset compared to the baseline period; this effect was significant on all training days. However, the magnitude of the increase in fluorescence significantly increased across days, with the smallest CS-related change in fluorescence occurring on day 1 and the largest CS-related change in fluorescence occurring on day 5 (**Figure 1G**, Two-way ANOVA, Training Day × Task Period p < 0.0001, Training Day p < 0.0001, Task Period p < 0.0001; Sidak’s Multiple Comparisons Test, Day 1 Baseline vs Day 3 Baseline p = 0.9949, Day 1 Baseline vs Day 5 Baseline p = 0.9684, Day 1 Baseline vs Day 1 CS p < 0.0001, Day 1 CS vs Day 3 CS p < 0.0001, Day 1 CS vs Day 5 CS p < 0.0001, Day 3 Baseline vs Day 5 Baseline p > 0.9999, Day 3 Baseline vs Day 3 CS p < 0.0001, Day 3 CS vs Day 5 CS p < 0.0001, Day 5 Baseline vs Day 5 CS p < 0.0001; N = 10 mice, n = 300 trials). There were no significant within-day differences in the amplitude of the dmPFC calcium signal when comparing dmPFC fluorescence during the 15 first trials to the last 15 trials within a given training day (**Supplemental Figure 2**). We also found no differences in calcium signal between successful and unsuccessful trials during the first second after CS onset; however, there were statistically significant differences during the later part of the PETH during the time window in which avoidance actions occur (**Supplemental Figure 3**). Taken together, these data suggest that there are learning-related increases in neural activity in the dmPFC during CS onset that become amplified across active avoidance learning.

**Figure 1.**
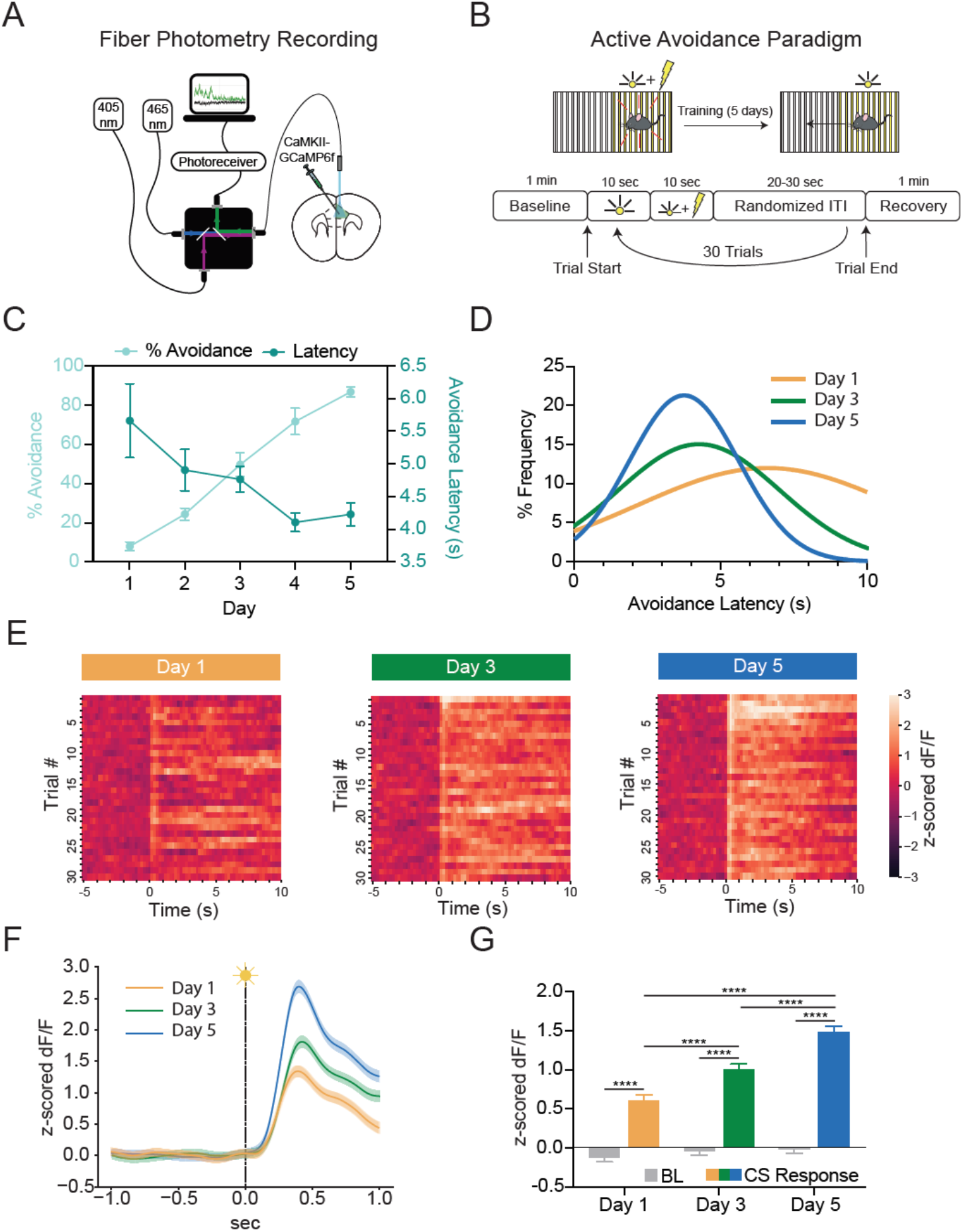
dmPFC shows learning-related increases in activity at CS onset during active avoidance learning. (A) Fiber photometry recording of dmPFC pyramidal neurons expressing GCaMP6f. (B) Behavioral schematic for active avoidance paradigm. (C) Average percent successful avoidance increased while avoidance latency decreased across training days. (D) Avoidance latency distribution shows avoidance latencies become shorter and more stereotyped across training. (E) Heatmaps of average change in calcium signal (z-scored dF/F) for each of the 30 trials presented in order from the first to the last trial for Day 1 (left), Day 3 (middle), and Day 5 (right). Heatmaps are aligned to CS onset (time zero) and show the total 10 second CS only period. dmPFC shows increased calcium signal at CS onset that becomes more consistent and sustained with training. (F) Perievent time histogram (PETH) showing increases in dmPFC calcium signal following CS onset. Orange line, mean ± standard error of the mean (SEM) for Day 1; green line, mean ± SEM for Day 3; blue line, mean ± SEM for Day 5. (G) Quantification of CS onset PETH shows calcium signal is significantly higher during the CS period (0 to 1 s) compared to the baseline period (−1 to 0 s) for all days. **** p ≤ 0.0001.

### dmPFC shows opposing patterns of activity during active avoidance and cued freezing

We next sought to examine dmPFC neural activity that corresponded to active avoidance and freezing behaviors that occurred later during the CS presentation. We investigated freezing behavior in addition to avoidance as freezing represents an alternative coping strategy that animals utilize early in learning before active coping strategies such as avoidance have been learned. The number of successful avoidances significantly increased across learning (**Figure 2A-B**, Repeated Measures One-way ANOVA p < 0.0001; Sidak’s Multiple Comparisons Test, Day 1 vs Day 3 p = 0.0002, Day 1 vs Day 5 p < 0.0001, Day 3 vs Day 5 p < 0.0001; N = 10 mice). When aligning the dmPFC calcium signal to avoidance onset on day 5 (**Figure 2C**), we found a statistically significant increase in fluorescence during the avoidance period compared to the baseline period (**Figure 2D**, Repeated Measures One-Way ANOVA p < 0.0001; Tukey’s Multiple Comparisons Test, Baseline vs Pre Avoid p < 0.0001, Baseline vs Avoid p < 0.0001, Baseline vs Post Avoid p < 0.0001, Pre Avoid vs Avoid p < 0.0001, Pre Avoid vs Post Avoid p = 0.6062, Avoid vs Post Avoid p < 0.0001; N = 10 mice, n = 253 trials). To rule out the possibility that these neural activity changes during avoidance onset in the dmPFC could be purely movement-related, we compared calcium signal during non-avoidance movements in the intertrial interval (ITI) period to avoidance movements of a similar velocity or duration from the same recording day. We found significantly increased fluorescence during avoidance movements compared to ITI movements during the time period where movements are initiated, suggesting that the increase in calcium signal during avoidance movements was not purely movement-related (**Supplemental Figure 4**). To further characterize the nature of the neural activity changes during avoidance, we created heatmaps of calcium activity on all individual trials aligned to avoidance onset and sorted them from shortest to longest avoidance latency (**Figure 2E**). In this heatmap, we observed a consistent time-locked peak in fluorescence that corresponded to avoidance onset. We also saw a sharp moving peak of fluorescence curving leftward that likely represented the increase in calcium signal at CS onset. These data suggest that the dmPFC separately encodes both the CS onset and avoidance onset through distinct increases in neural activity.

**Figure 2.**
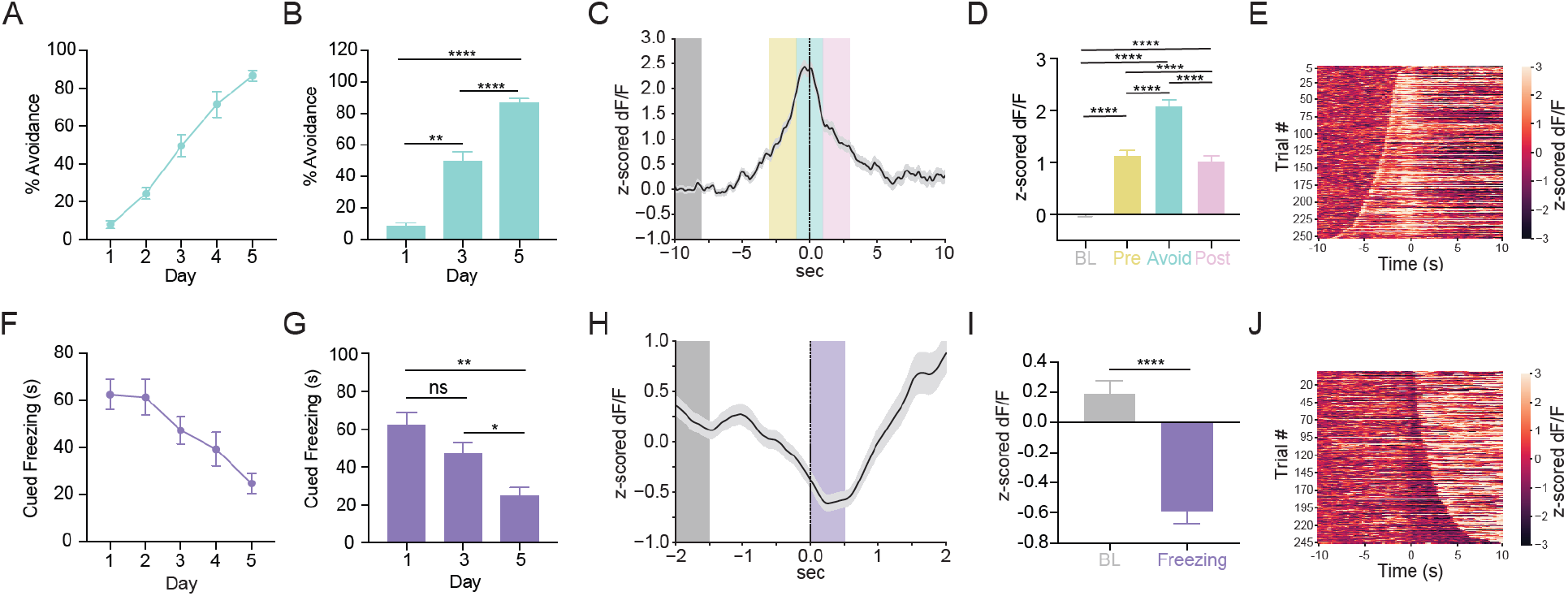
dmPFC shows opposing patterns of activity during active avoidance and cued freezing behavior. (A) Percent successful avoidance across training days. (B) Quantification of percent successful avoidance shows animals significantly increase avoidance across training. (C) PETH shows an increase in calcium signal at avoidance onset. Line with shading represents mean ± SEM. Grey box, baseline period (BL); yellow box, pre avoidance period (Pre); teal box, avoidance period (Avoid); pink box, post avoidance period (Post). (D) Quantification of avoidance PETH reveals significantly increased calcium signal in pre avoid (−3 to −1 s), avoid (−1 to 1 s), and post avoid (1 to 3 s) periods compared to the baseline period (−10 to −8 s). (E) Heatmap of change in calcium signal for individual avoidance trials aligned to avoidance onset and sorted from shortest to longest avoidance latency. Heatmap shows distinct increases in calcium signal at CS onset (slope curving leftward) and avoidance onset (time zero). (F) Percent freezing during the CS only period (cued freezing) across training days. (G) Quantification of percent cued freezing shows that animals significantly decrease cued freezing across training. (H) PETH shows decrease in calcium signal at freezing onset. Line with shading represents mean ± SEM. Grey box, baseline period (BL); Purple box, freezing period (Freezing). (I) Quantification of freezing PETH shows significant decrease in calcium signal during the freezing period (0-0.5 s) compared to the baseline period (−2 to −1.5 s). (J) Heatmap of change in calcium signal during individual freezing bouts aligned to freezing onset and sorted from shortest to longest freezing bout. Heatmap shows dips in calcium signal at freezing onset that increases in length as freezing bout duration increases. ns = not significant, * p ≤ 0.05, ** p ≤ 0.01, **** p ≤ 0.0001.

In contrast to avoidance, the amount of freezing during the CS-only period (cued freezing) significantly decreased across learning (**Figure 2F-G**, Repeated Measures One-way ANOVA p = 0.0045; Sidak’s Multiple Comparisons Test, Day 1 vs Day 3 p = 0.4807, Day 1 vs Day 5 p = 0.0024, Day 3 vs Day 5 p = 0.023; N = 10 mice). When we generated a PETH of dmPFC calcium activity aligned to freezing onset on day 1 for all cued freezing bouts with a 1 second minimum duration (**Figure 2H**), we found a statistically significant decrease in fluorescence during the freezing period compared to the baseline period (**Figure 2I**, Paired t-test p < 0.0001; N = 10 mice, n = 246 trials). When examining a heatmap of calcium activity on all individual trials aligned to freezing onset and sorted by shortest to longest freezing bout duration, we saw a dip in fluorescence at freezing onset that increased in duration with longer freezing bouts (**Figure 2J**). This suggested that the duration of the decrease in dmPFC calcium activity during freezing corresponded to the duration of the freezing bout length, providing further evidence that the dip in fluorescence was tightly time-locked with freezing behavior. Overall, our results suggest that the dmPFC shows opposing patterns of activity during avoidance and freezing and that these patterns of activity are distinct from the neural activity observed during CS onset.

### dmPFC-DMS and dmPFC-BLA projections show learning-related increases in activity at CS onset

We also explored how subpopulations of dmPFC neurons defined by their downstream projection target may diverge in the encoding of active avoidance. To obtain projection-specific fiber photometry recordings from the dmPFC-DMS projection, we used a dual virus retrograde targeting strategy to express GCaMP only in dmPFC neurons projecting to the DMS (**Figure 3A, Supplemental Figure 5**). Behavioral results from this cohort revealed that the mice learned to 80% successful avoidance by day 5, average avoidance latencies were between 4-6 seconds, and avoidance latency decreased across training (**Figure 3B**). To visualize potential task-relevant information within the dmPFC-DMS projection, we examined heatmaps of the average calcium signal change on each trial for the first 10 seconds of the CS (CS-only period) (**Figure 3C**). In the dmPFC-DMS projection, already on the first day of learning we saw a sustained increase in fluorescence during the CS only period, although the start of the signal did not appear clearly time locked to CS onset and the sustained increase did not appear on every trial. However, by day 5 of learning the sustained increase in fluorescence in the dmPFC-DMS projection became time locked to CS onset and consistently appeared on every trial. When examining calcium activity during the first second of CS onset in the dmPFC-DMS projection (**Figure 3D**), we found a significant increase in signal at CS onset compared to baseline on day 5 but not on day 1. We additionally found that there was a significant difference in calcium signal at CS onset across days, with a larger CS-evoked increase in signal on day 5 compared to day 1, suggesting that there were learning-related changes (**Figure 3E**, Two-way ANOVA, Training Day × Task Period p = 0.0498, Training Day p = 0.0725, Task Period p < 0.0001; Sidak’s Multiple Comparisons Test, Day 1 Baseline vs Day 1 CS p = 0.0634, Day 1 Baseline vs Day 5 Baseline p > 0.9999, Day 1 CS vs Day 5 CS p = 0.0466, Day 5 Baseline vs Day 5 CS p < 0.0001; N = 8 mice, n = 300 trials).

**Figure 3.**
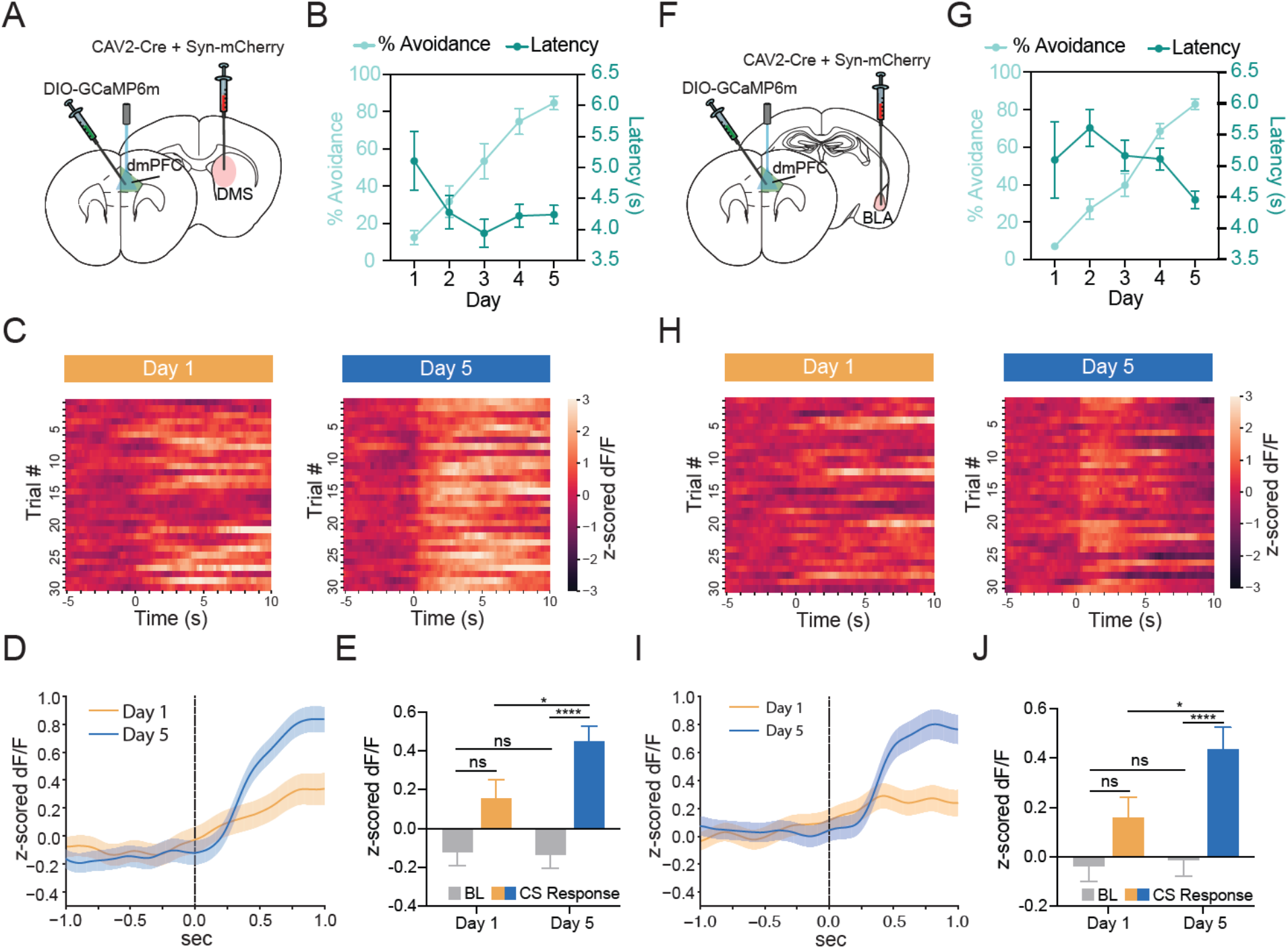
dmPFC-DMS and dmPFC-BLA show similar learning-related increases in activity at CS onset during active avoidance learning. (A) Viral targeting strategy for dmPFC-DMS photometry. (B) Percent avoidance increases while avoidance latency decreases across training in the dmPFC-DMS cohort. (C) Heatmaps of change in calcium signal aligned to CS onset for each of the 30 trials arranged from first to the last trial for Day 1 (left) and Day 5 (right). dmPFC-DMS projection shows sustained increases in calcium signal at CS onset that become more consistent across training. (D) PETH shows increases in signal at CS onset in the dmPFC-DMS projection following training. orange line, mean ± SEM for Day 1; blue line, mean ± SEM for Day 5. (E) Quantification of the dmPFC-DMS CS onset PETH shows significant increase in calcium signal during the CS period (0 to 1 s) compared to the baseline period (−1 to 0 s) for Day 5, but not Day 1. (F) Viral targeting strategy for dmPFC-BLA photometry. (G) Percent avoidance increases while avoidance latency decreases across training in the dmPFC-BLA cohort. (H) Heatmaps of change in calcium signal aligned to CS onset for each of the 30 trials arranged from first to the last trial for Day 1(left) and Day 5 (right). dmPFC-BLA projection shows transient increases in calcium signal at CS onset only during later stages of training. (I) PETH shows increases in signal at CS onset in the dmPFC-BLA projection following training. orange line, mean ± SEM for Day 1; blue line, mean ± SEM for Day 5. (J) Quantification of the dmPFC-BLA CS onset PETH shows significant increase in calcium signal during the CS period (0 to 1 s) compared to the baseline period (−1 to 0 s) for Day 5, but not Day 1. ns = not significant, * p ≤ 0.05, **** p ≤ 0.0001.

We next examined neural activity in the dmPFC-BLA projection during active avoidance learning using the same dual virus retrograde targeting strategy (**Figure 3F, Supplemental Figure 5**). Behaviorally, we saw similar trends to the dmPFC-DMS projection cohort (**Figure 3G**). Heatmaps of the average calcium activity change during the first 10 seconds of the CS revealed that the dmPFC-BLA projection did not show clearly organized patterns of fluorescence on the first day of learning. However, by day 5 this projection showed a clear transient increase in fluorescence that was time locked to CS onset and consistently seen across trials (**Figure 3H**). When examining calcium activity in the dmPFC-BLA during the first second of CS onset across learning (**Figure 3I**), the dmPFC-BLA projection showed no significant differences in signal between the baseline period and CS onset on day 1, but showed significant increases in calcium signal at CS onset compared to the baseline period on day 5. We also found that there was a significant increase in signal at CS onset on day 5 of learning compared to day 1 of learning (**Figure 3J**, Two-way ANOVA, Training Day x Task Period p = 0.0816, Training Day p = 0.0411, Task Period p < 0.0001; Sidak’s Multiple Comparisons Test, Day 1 Baseline vs Day 5 Baseline p > 0.9999, Day 1 Baseline vs Day 1 CS p = 0.3023, Day 1 CS vs Day 5 CS p = 0.0442, Day 5 Baseline vs Day 5 CS p < 0.0001; N = 9 mice, n = 300 trials). Additional analyses examining calcium activity in these projections during successful and unsuccessful trials found that the CS-evoked fluorescence changes during successful trials did not significantly differ from that on unsuccessful trials for either projection (**Supplemental Figure 6**). Overall, our results suggest that both the dmPFC-DMS and dmPFC-BLA projections show learning-related increases in neural activity at CS onset during active avoidance learning.

### dmPFC-DMS and dmPFC-BLA projections show divergent encoding of active avoidance behavior

We were additionally interested in examining projection-specific neural activity during avoidance and freezing behaviors. Both cohorts reached 80% successful avoidance by day 5 of learning (**Figure 4A-C**, dmPFC-DMS Paired t-test p < 0.0001, dmPFC-BLA Paired t-test p < 0.0001; dmPFC-DMS N = 8 mice, dmPFC-BLA N = 9 mice). While calcium activity in these two projections was similar upon CS onset, we found a striking contrast in avoidance-related calcium activity between the dmPFC-DMS and dmPFC-BLA projections. In the PETH aligned to avoidance onset, while the dmPFC-DMS projection showed a hill-like increase in fluorescence at avoidance onset, the dmPFC-BLA projection showed a descending slope (**Figure 4D**). Validating these stark changes, the dmPFC-DMS projection showed a significant increase in signal during the avoidance period compared to the baseline period while the dmPFC-BLA projection showed a significant decrease in signal between the pre-avoidance and post avoidance periods. In addition, the dmPFC-DMS and the dmPFC-BLA calcium signals were distinct from each other as they statistically differed throughout the avoidance and post-avoidance periods (**Figure 4E**, Two-way ANOVA, Task Period × Projection p < 0.0001, Task Period p < 0.0001, Projection p < 0.0001; Sidak’s Multiple Comparisons Test, dmPFC-DMS Baseline vs dmPFC-DMS Avoid p < 0.0001, dmPFC-BLA Pre Avoid vs dmPFC-BLA Post Avoid p < 0.0001, dmPFC-DMS Baseline vs dmPFC-BLA Baseline p > 0.9999, dmPFC-DMS Pre Avoid vs dmPFC-BLA Pre Avoid p = 0.9837, dmPFC-DMS Avoid vs dmPFC-BLA Avoid p < 0.0001, dmPFC-DMS Post Avoid vs dmPFC-BLA Post Avoid p < 0.0001; dmPFC-DMS N = 8 mice, n = 195 trials, dmPFC-BLA N = 9 mice, n = 211 trials). Using movements of similar velocity or duration during the ITI period as a control, we found significant differences in fluorescence between the ITI movements compared to avoidance movements, suggesting that the changes in calcium activity in these projections during avoidance onset were not purely movement-related (**Supplemental Figure 7**). To further characterize the avoidance-related activity seen in these projections, we created heatmaps of calcium activity on all individual trials aligned to avoidance onset sorted from shortest to longest avoidance latency for each projection (**Figure 4F**). In the dmPFC-DMS projection avoidance heatmap, we saw an increase in fluorescence that curved leftwards, which corresponded to the start of the CS. This increased fluorescence that occurred at CS onset was sustained through avoidance onset as there were no clear distinctions in signal between when the CS began and when the avoidance began. In contrast, in the dmPFC-BLA projection heatmap, CS onset and avoidance onset were marked by distinct changes in calcium activity. There was a clear increase in fluorescence sloping leftward that corresponded to CS onset, whereas avoidance onset was marked by a time-locked drop in fluorescence.

**Figure 4.**
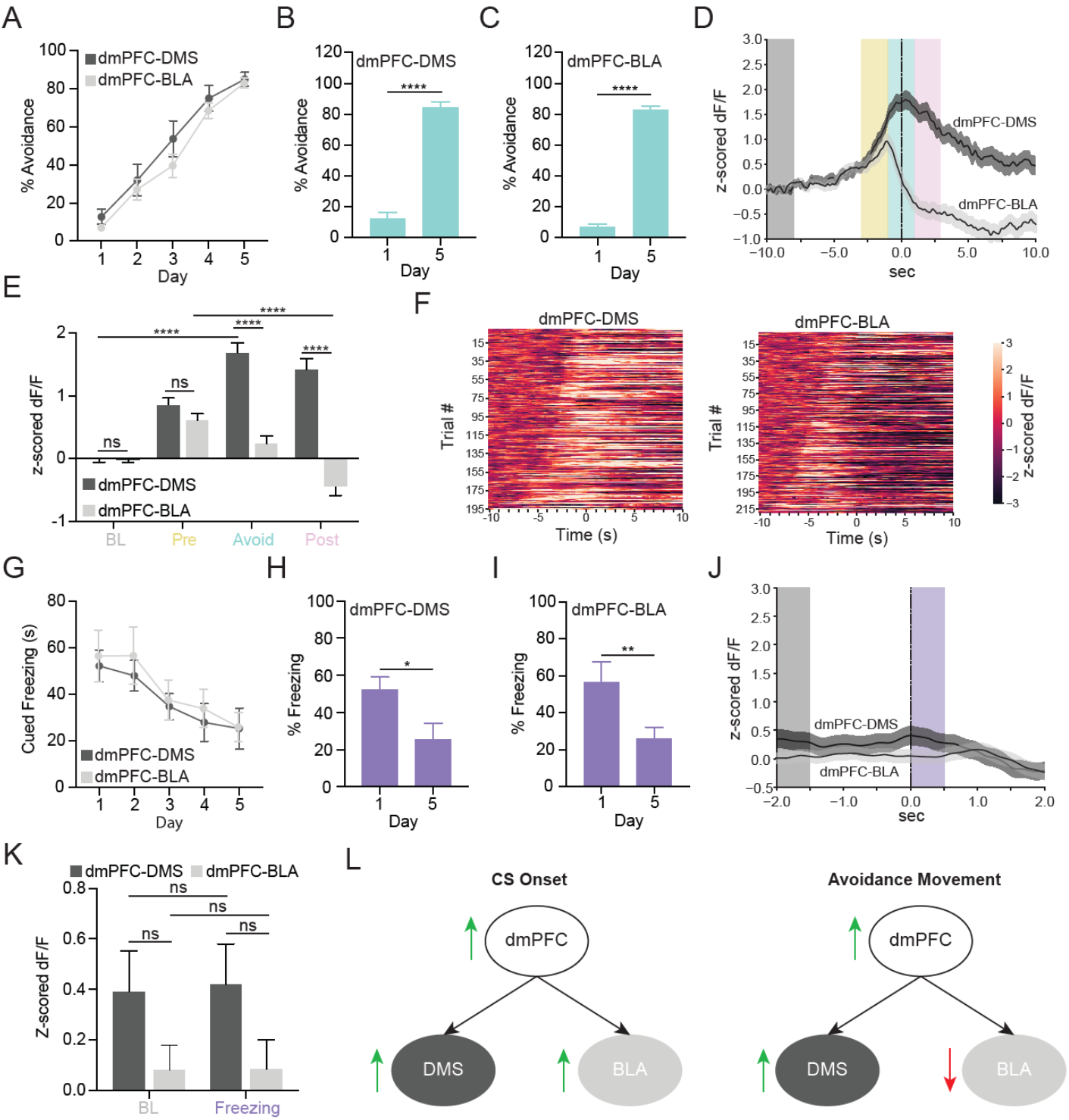
dmPFC-DMS and dmPFC-BLA show divergent encoding of active avoidance behavior. (A) Percent avoidance across training days in the dmPFC-DMS (dark grey line) and dmPFC-BLA (light grey line) cohort. (B-C) Percent avoidance significantly increases from Day 1 to Day 5 in the dmPFC-DMS (left) and dmPFC-BLA (right) cohort. (D) PETH shows increase in calcium signal in the dmPFC-DMS projection and decrease in calcium signal in the dmPFC-BLA projection during avoidance onset. Dark grey line, mean ± SEM for dmPFC-DMS projection; light grey line, mean ± SEM for dmPFC-BLA projection; Grey box, baseline period (BL); yellow box, pre avoidance period (Pre); teal box, avoidance period (Avoid); pink box, post avoidance period (Post). (E) Quantification of avoidance PETH shows a significant increase in calcium signal in the avoid (−1 to 1 s) period compared to baseline period (−10 to −8 s) for dmPFC-DMS projection and a significance decrease in signal during the post avoid (1 to 3 s) period compared to the pre avoid (−3 to −1 s) period in the dmPFC-BLA projection. (F) Heatmap of change in calcium signal for individual avoidance trials aligned to avoidance onset and sorted from shortest to longest avoidance latency for the dmPFC-DMS (left) and dmPFC-BLA (right) projections. (G) Percent cued freezing in the dmPFC-DMS (dark grey line) and the dmPFC-BLA (light grey line) cohort. (H-I) Percent cued freezing significantly decreases from Day 1 to Day 5 in the dmPFC-DMS (left) and the dmPFC-BLA (right) cohorts. (J) PETH shows no change in calcium signal at freezing onset for either the dmPFC-DMS or the dmPFC-BLA projection. Dark grey line, mean ± SEM for dmPFC-DMS projection; light grey line, mean ± SEM for dmPFC-BLA projection; Grey box, baseline period (BL); Purple box, freezing period (Freezing). (K) Quantification of freezing PETH shows no significant change in calcium signal during the freezing period (0-0.5 s) compared to the baseline period (−2 to −1.5 s). (L) Graphical abstract summarizing main findings from the study. ns = not significant * p ≤ 0.05, ** p ≤ 0.01, **** p ≤ 0.0001.

We next examined how the dmPFC-DMS and the dmPFC-BLA projections encoded freezing behavior and found statistically significant decreases in freezing on day 5 compared to day 1 for each projection (**Figure 4G-I**, dmPFC-DMS Paired t-test p = 0.0484, dmPFC-BLA Paired t-test p = 0.0032; dmPFC-DMS N = 8 mice, dmPFC-BLA N = 9 mice). However, there was no significant difference in signal between the baseline period and the freezing period in the perievent time histograms aligned to freezing onset for each projection (**Figure 4J-K**, Two-way ANOVA, Task Period × Projection p = 0.9234, Task Period p = 0.8965, Projection p = 0.0145; Sidak’s Multiple Comparisons Test, dmPFC-DMS Baseline vs dmPFC-BLA Baseline p = 0.4562, dmPFC-DMS Baseline vs dmPFC-DMS Freezing p > 0.9999, dmPFC-BLA Baseline vs dmPFC-BLA Freezing p > 0.9999, dmPFC-DMS Freezing vs dmPFC-BLA Freezing 0.3624; dmPFC-DMS N = 8 mice, n = 183 trials, dmPFC-BLA N = 9 mice, n = 229 trials). Overall, our results show opposing patterns of activity in the dmPFC-DMS and dmPFC-BLA projection during active avoidance behavior, with increased activity in the dmPFC-DMS projection and decreased activity in the dmPFC-BLA projection at avoidance onset. The main findings from our study are summarized in **Figure 4L**.

## DISCUSSION

We found that the dmPFC and its projections to the DMS and the BLA contain learning-related increases in activity at CS onset during active avoidance. Encoding of active avoidance diverged in the dmPFC-DMS and dmPFC-BLA projections, which showed increased and decreased neural activity at avoidance onset, respectively. To our knowledge, this is the first study to record the endogenous activity of distinct dmPFC projections during active avoidance behavior. Our results reveal the importance of studying projection-defined dmPFC subpopulations as they may play distinct but complementary roles in active avoidance learning and expression.

The sharp peak of dmPFC activity at CS onset that significantly increased in amplitude across training suggests that the dmPFC encodes learning-related information for active avoidance behavior. Given that significant differences in neural activity were seen across days but not within days suggests that the learning-related increase in activity at CS onset in the dmPFC is a consolidated phenomenon that gradually builds across time. Another recent study used dmPFC single unit activity to successfully decode CS identity between a CS that predicted shock and led to avoidance (CS+) and a control CS that did not predict shock and did not lead to avoidance (CS-) (Jercog et al., 2021), corroborating our finding that the dmPFC holds active avoidance task-relevant information. While our results show increased dmPFC activity aligned with CS onset, another study in rats using a platform-mediated active avoidance task found inhibition of single dmPFC units upon CS onset unique to avoidance (Diehl et al., 2018). This discrepancy could be explained by the subregion of dmPFC targeted (rostral vs caudal), or technical differences between bulk calcium recording and single unit electrophysiology. For example, calcium indicators are more sensitive to increases rather than decreases in activity and may preferentially reflect synchronous and/or bursting activity of groups of neurons (Chen et al., 2013). Interestingly, there was no difference in this CS-evoked neural signal between successful and unsuccessful trials. This observation is supported by other studies (Diehl et al., 2018; Jercog et al., 2021) and suggests that this activity may signal the option to avoid rather than the avoidance behavior itself. Of note, the initial sharp peak of activity upon CS onset was present on the first day before learning had occurred, albeit significantly smaller in amplitude than on the last day of training. Given that the dmPFC receives various sensory related inputs (Ährlund-Richter et al., 2019), this initial peak on day 1 may represent sensory features of the CS, while the increase in amplitude of this peak across days is reflective of learning-related activity. Overall, this is the first study to our knowledge to examine longitudinal learning-related changes in dmPFC activity across days of an active avoidance task.

The dmPFC also showed a robust increase in neural activity during avoidance onset in our task. This result is consistent with a recent study employing the platform-mediated avoidance task, which also found increased activity in the dmPFC when animals moved onto a platform to avoid shock. However, there was no difference in the proportion of cells excited between mice trained on fear conditioning or active avoidance in the same apparatus (Diehl et al., 2018), suggesting that increased activity was not specific to avoidance behavior in their task. While their study controlled for locomotion by comparing platform entries between separate avoidance-trained and fear-conditioned cohorts, here we performed a within-animal locomotor control. Comparing dmPFC neural activity during avoidance movements versus intertrial interval movements of similar duration and velocity, we found that the increased neural activity seen during avoidance is not accounted for by general movement alone. This finding is corroborated by another study using dmPFC activity to decode avoidance behavior in a discriminative two-way active avoidance task, which found an increase in decoding accuracy within the last second before the avoidance movement which could not be accounted for by speed (Jercog et al., 2021). The predictive increase in decoder accuracy before avoidance initiation is also in alignment with the increase in activity we observed in the dmPFC preceding avoidance onset. Furthermore, only the excitatory responses in the dmPFC contained predictive information about avoidance initiation in the discriminative two-way active avoidance task (Jercog et al., 2021), which further supports that notion that the excitatory dmPFC activity we see in our task contains crucial information for proper avoidance performance rather than only encoding movement. We also found differences in dmPFC activity between successful and unsuccessful trials during the period where avoidances normally occur, which has been similarly identified in other studies and may correspond to differences in the behavioral repertoire of the animals during successful and unsuccessful trials (Diehl et al., 2018; Jercog et al., 2021).

We propose that the increased dmPFC activity at avoidance onset may be important for the animal to take action in the face of an anxiogenic stimulus. In the active avoidance task, when the CS light is on, dmPFC activity increases when the animal initiates an avoidance movement within the anxiogenic lit chamber of the apparatus. A recent study from our laboratory using the elevated zero maze to assess approach-avoidance conflict showed that dmPFC activity increases as the animal moves into the anxiogenic open arms of the maze (Loewke et al., 2021). These seemingly disparate findings may be reconciled by the idea that dmPFC activity allows the animal to explore or take action in the face of an anxiogenic stimulus, while dmPFC activity decreases once the anxiogenic stimulus has been successfully avoided (i.e., shuttling to the safe chamber in active avoidance, and entering the closed arm of the elevated zero maze). The notion that dmPFC activity may be important for resolving conflicting signals between the drive to explore or take action and the drive to passively cope with an anxiogenic stimulus is supported by various studies suggesting a role for the dmPFC in decision making under conflict (Burgos-Robles et al., 2017; Friedman et al., 2015; Ishikawa et al., 2020; Loewke et al., 2021).

The dmPFC as a whole showed decreased activity during freezing in our active avoidance task, with the duration of this decrease in activity corresponding to the freezing bout length. In contrast, in vivo electrophysiology studies have found increased firing rates in dmPFC neurons during freezing behavior in classical and discriminative fear conditioning tasks (Burgos-Robles et al., 2009; Dejean et al., 2016; Likhtik et al., 2014). Given that calcium indicators are more sensitive to increases rather than decreases in activity (Chen et al., 2013), this difference seems likely unrelated to technique used and may instead be due to key differences in the tasks, such as the fact that the active avoidance task allows for both passive and active coping responses to threat, whereas in classical fear conditioning animals have no control over the shocks and therefore are biased toward passive coping via freezing. Future studies using single cell resolution calcium imaging will help elucidate the encoding of individual dmPFC neurons during freezing in active avoidance versus fear conditioning tasks.

Both the dmPFC-DMS and dmPFC-BLA projections showed increased activity at CS onset, with learning-related changes evidenced by significant increases in signal amplitude across training in both projections. As the dmPFC-DMS projection plays an important role in goal-directed behavior (Hart, Bradfield, & Balleine, 2018; Hart, Bradfield, Fok, et al., 2018), this CS-related activity could hold crucial information regarding action-outcome contingencies for this task. The dmPFC-BLA projection has been linked to associative fear conditioning (Adhikari et al., 2015; Cho et al., 2013) and thus CS-related activity in this projection may contain key information on CS-US associations in this task. When comparing successful and unsuccessful trials, we found no differences in activity during CS onset in either projection, suggesting that CS-related activity in these projections may again signal an avoidance option rather than avoidance behavior itself. Interestingly, downstream BLA neurons do show distinct activity on successful and unsuccessful avoidance trials (Kyriazi et al., 2018). Thus, the BLA likely receives information necessary for distinguishing between these trial types from a region outside the dmPFC. Future studies should attempt to uncover additional circuits that differentiate between successful and unsuccessful trials that may act upstream of the BLA.

While CS-aligned activity looked similar in both projections, they displayed opposing patterns of activity at avoidance onset, with the dmPFC-DMS projection showing increased activity and the dmPFC-BLA projection showing decreased activity. The dmPFC-DMS projection directly interfaces downstream with the striatum which regulates motor control and action selection (Kravitz & Kreitzer, 2012) and is therefore poised to play a privileged role in aiding avoidance movement initiation. The striatum consists of D1 and D2 medium spiny neurons (MSNs) that, when optogenetically stimulated, drive motor initiation and motor cessation, respectively (Kravitz & Kreitzer, 2012; Redgrave et al., 2010). The mPFC has stronger synaptic input to D1 versus D2 MSNs, and optogenetic stimulation of D1 MSNs recapitulates anxiolytic effects seen with dmPFC-DMS stimulation (Loewke et al., 2021). Increased activity in the dmPFC-DMS projection may directly excite striatal D1 MSNs leading to motor initiation and, in our task, active avoidance behavior. Conversely, the dmPFC-BLA projection has been tied to freezing behavior, with dmPFC-BLA stimulation during fear conditioning leading to increased freezing at extinction recall (Adhikari et al., 2015). Fear-related information is thought to be sent from the BLA to the central amygdala (CeA) to downstream brainstem structures leading to freezing initiation (Tovote et al., 2015). Given that increased activity in the dmPFC-BLA-CeA pathway may promote freezing, the decreased activity we see in the dmPFC-BLA projection during avoidance behavior may help suppress freezing to allow proper active avoidance behavior to occur. The contrasting neural activity in the dmPFC-DMS and dmPFC-BLA projections may therefore play distinct yet complementary roles in coordinating successful active avoidance behavior through the initiation of avoidance movements (dmPFC-DMS) and the suppression of freezing behavior (dmPFC-BLA).

While the dmPFC-DMS projection has not been previously explored within the context of active avoidance, a recent optogenetic study has causally implicated the dmPFC-BLA projection in platform-mediated active avoidance (Diehl et al., 2020). Stimulation of the dmPFC-BLA projection increases avoidance in the platform-mediated task (Diehl et al., 2020), while our photometry results would suggest that inhibiting the dmPFC-BLA projection may increase avoidance given that dmPFC-BLA activity decreases acutely during avoidance in our task. In our previous study examining the dmPFC-DMS and dmPFC-BLA projections during an innate approach-avoidance task, we found that the dmPFC-DMS projection recapitulated whole population dmPFC activity while the dmPFC-BLA projection did not (Loewke et al., 2021). Similarly, here we find that the dmPFC-DMS projection shows increased activity during avoidance similar to the dmPFC overall, while the dmPFC-BLA projection shows distinct decreases in activity during avoidance. The projection-specific activity we observed during avoidance intriguingly parallels fMRI findings during active avoidance in humans (Collins et al., 2014; Delgado et al., 2009). In one study, coupling between the mPFC and the caudate (the human equivalent of the DMS) and between the mPFC and the amygdala during active avoidance trials predicted better active avoidance performance (Collins et al., 2014). The increased coupling between mPFC and caudate/amygdala during active avoidance performance parallels the signals we see in the dmPFC-DMS and dmPFC-BLA projections during active avoidance behavior. The human study also found increased activity in the caudate and decreased activity in the amygdala during active avoidance behavior (Collins et al., 2014), similar to the increased activity in the dmPFC-DMS projection and the decreased activity in the dmPFC-BLA projection we observed during active avoidance. Overall, these results highlight the importance of the mPFC downstream communication with both the dorsal striatum and the amygdala and suggest conservation of function across species in these circuits during active avoidance behavior.

Overall, we find task-relevant information encoding in the dmPFC and its projections to the DMS and the BLA during active avoidance learning, with opposing patterns of activity in the dmPFC-DMS and dmPFC-BLA projections during active avoidance behavior, suggesting that these circuits play distinct but complementary roles in the successful enactment of active avoidance behavior. These findings provide a crucial first step in identifying precise prefrontal subpopulations and circuits for active avoidance behavior that may help guide future treatment targets to alleviate avoidance symptoms seen in anxiety disorders.

## MATERIALS AND METHODS

### Animals

We used wild-type C57BL6/J mice purchased from Jackson Laboratories. Animals were raised in normal light conditions (12:12 light/dark cycle) and given food and water ad libitum. All experiments were conducted in accordance with procedures established by the Institutional Animal Care and Use Committee at the University of California, San Francisco.

### Stereotaxic Surgery, Viral Injections, and Fiber Optic Cannula Implantation

Surgeries were performed at 10-14 weeks of age. Mice were anesthetized using 5.0% isoflurane at an oxygen flow rate of 1 L/min and placed on top of a heating pad in a stereotaxic apparatus (Kopf Instruments, Tujunga, CA, USA). Anesthesia was maintained with 1.5-2.0% isoflurane for the duration of the surgery. Respiration and toe pinch response were monitored closely. Slow-release buprenorphine (0.5 mg/kg) and ketoprophen (1.6 mg/kg) were administered subcutaneously at the start of surgery. The incision area was shaved and cleaned with ethanol and betadine. Lidocane (0.5%) was administered topically on the scalp. An incision was made along the midline and bregma was measured. Virus was injected (as described below) using a 10 μL nanofil syringe (World Precision Instruments, Sarasota, FL, USA) with a 33-gauge beveled needle. We used an injection rate of 100 nL/min with a 10-minute delay before retracting the needle. Mice recovered in a clean cage on top of a heating pad and a subsequent injection of ketoprofen (1.6 mg/kg) was given the following day.

For fiber photometry, we injected 500 nL of AAV5-CaMKII-GCaMP6f or AAV5-CaMKII-eYFP into the dmPFC to record pyramidal neuron activity; to record dmPFC-DMS and dmPFC-BLA projection neurons, we injected 1500 nL of AAV1-Syn-Flex-GCaMP6m or AAV5-EF1a-DIO-eYFP into the dmPFC and 500 nL of CAV2-Cre and hSyn-mCherry into the DMS and BLA. Injection coordinates (in millimeters relative to bregma) were as follows: dmPFC (1.8 A/P, −.35 M/L, −2.4 D/V), DMS (.8 A/P, −1.5 M/L, −3.5 D/V), BLA (−1.4 A/P, −3.3 M/L, −4.9 D/V). For all fiber photometry experiments, we implanted a 2.5 mm metal fiber optic cannula with 400 μm fiber optic stub (Doric Lenses, Quebec, Canada) in the dmPFC and waited 4-5 weeks for viral expression. Implant coordinates for the mPFC were 1.8 A/P, −.35 M/L, −2.2 D/V.

All viruses were obtained from Addgene, UNC Vector Core, or Institut de Génétique Moléculaire de Montpellier, Montpellier, France.

### Active Avoidance Behavior

Mice underwent a two-way active avoidance procedure adapted from Pare 2018. Active avoidance training occurred in a custom made apparatus consisting of two shock floors with strips of visible spectrum LED lights underneath each shock floor. Both shock and light presentations were controlled by an arduino using custom-made arduino code (Arduino, Somerville, MA, USA) in conjunction with location data from video recording software, Ethovision XT (Noldus, Wageningen, Netherlands). All trials were conducted in the dark and infrared lights beneath each shock floor were used to track the animals. Mice underwent 30 active avoidance trials per day for 5 days. Each active avoidance trial consisted of a 10 second light cue followed by 10 seconds of light plus 0.3 mA shock. Light and shock were presented on the shock floor the mouse was currently on at the initiation of the trial. Mice were able to avoid the shock altogether by moving onto the other unlit shock floor during the 10 second light only period. This was considered a successful active avoidance trial. Trials in which the mouse failed to move to the other unlit shock floor during the 10 seconds of light only are considered unsuccessful trials. Training continued until the group average was at or above 80% successful avoidance (24 out of 30 trials). Location of the mice was recorded and quantified using Ethovision XT software.

### Fiber Photometry Recording

*In vivo* calcium data were acquired using a custom-built rig based on a previously described setup (Lerner et al., 2015). This setup was controlled by an RZ5P fiber photometry processor (TDT, Alachua, FL, USA) and Synapse software (TDT). The RZ5P/Synapse software controlled a 4 channel LED Driver (DC4100, Thorlabs, Newton, NJ, USA) which in turn controlled two fiber-coupled LEDS: 470 nm for GCaMP stimulation and 405 nm to control for artifactual fluorescence (M470F3, M405FP1, Thorlabs). These LEDs were sinusoidally modulated at 210 Hz (470 nm) and 320 Hz (405 nm) and connected to a Fluorescence Mini Cube with 4 ports (Doric Lenses) and the combined LEF output was connected through a fiber optic patch cord (0.48 NA, 400 μm, Doric Lenses) to the cannula via a ceramic sleeve (Thorlabs). The emitted light was focused onto a Visible Femtowatt Photoreceiver Module (Model 2151, Newport, AC low) and sampled at 60 Hz. Video tracking software (Ethovision, Noldus) was synchronized to the photometry setup using TTL pulses generated every 10 seconds following the start of the Noldus trial. Raw photoreceiver data was extracted and analyzed using custom scripts in Matlab (The MathWorks, Natick, MA, USA). The two output signal data was demodulated from the raw signal based on the LED modulation frequency. To normalize the data and correct for bleaching, the 405 nm channel signal was fitted to a polynomial over time and subtracted from the 470 nm GCaMP signal, yielding the DF/F value.

### Perfusions and Histology

Following the conclusion of behavioral experiments, animals were anesthetized using 5% isoflurane and given a lethal dose (1.0 mL) cocktail of ketamine/xylazine (10 mg/ml ketamine, 1 mg/ml xylazine). They were then transcardially perfused with 10 mL of 1X PBS followed by 10 mL 4% paraformaldehyde (PFA). Brains were extracted and left in 4% PFA overnight and then transferred to a 30% sucrose solution until slicing. The brains were frozen and sliced on a sliding microtome (Leica Biosystems, Wetzlar, Germany) and placed in cryoprotectant in a well-plate. Slices were then washed in 1X PBS, mounted on slides (Fisherbrand Superfrost Plus, ThermoFisher Scientific, Waltham, MA, USA) and air dried (covered). ProLong Gold antifade reagent (Invitrogen, ThermoFisher Scientific) was injected on top of the slices and a cover slip (Slip-rite, ThermoFisher Scientific) was placed on top and the slides were left to dry overnight (covered). Viral injection, fiber photometry cannula implant, and optogenetic cannula implant placements were histologically verified on a fluorescence microscope (Leitz DMRB, Leica).

### Movement and Freezing Behavior Analysis

Following the recording of location data using Ethovision, post data collection analysis was performed to identify movement initiations using Ethovision’s built in movement detection software. The detection settings used were a 10 sample averaging window, 2.25 cm/sec start velocity threshold, and 2 cm/sec stop velocity threshold. Additionally, we used open source code (Pennington et al., 2019) to identify freezing. The parameters we used for this analysis were a motion cuttoff of 9.0, freezing threshold of 1000, and minimum freeze duration of 25 samples (1 second).

### Fiber Photometry Data Analysis

Data was analyzed in PyCharm CE (JetBrains, Prague, Czechia) environment. Behavioral, location, and movement initiation data was extracted from both Ethovision and Arduino and synced to Synapse fiber photometry data. From this we extracted the behavioral data (percent avoidance, avoidance latency, and freezing) across all five days of learning. Additionally, we generated peri-event time histograms and heatmaps by time-locking the neural activity (dF/F) and z-scoring the signal to the baseline period (last 10 seconds of inter-trial-interval (ITI) preceding the event). These events included CS (light) onset (also split into successful and unsuccessful trials), avoidance movement initiation (movements during the 10 second light only period of successful trials), and freezing behavior initiation (freezing during the 10 second light only period of all trials). In addition, we also analyzed movement initiations during the ITI periods across all days. The heatmaps for avoidance movements and freezing were sorted by avoidance latency and freezing duration respectively. Quantification was done using the average signal across the following time windows:

CS onset: Baseline (−1 to 0 sec), CS response (0 to 1 sec)
CS successful vs. unsuccessful: Baseline (−1 to 0 sec), Initial CS response (0 to 1 sec), Pre-avoidance (1 to 2 sec), Post-avoidance (9 to 10 sec)
Avoidance movement: Baseline (−10 to −8 sec), Pre-avoidance (−3 to −1 sec), Avoidance (−1 to 1 sec), Post-avoid: (1 to 3 sec)
ITI movement: Baseline (−10 to −8 sec), Pre-movement (−3 to −1 sec), Movement (−1 to 1 sec), Post-movement: (1 to 3 sec)
Freezing: Baseline (−2 to −1.5 sec), Freezing (0 to 0.5 sec)

All other non-avoidance movement controls were quantified identically to avoidance movement. Lastly, histograms of the distribution of velocity and movement duration for all movement parameters were generated in Prism using a bin width of 1 cm/sec and 1 second respectively.

### Statistical Analysis

Statistical Analysis was performed with Prism 8 (Graphpad Software, San Diego, CA, USA). Normality was tested with D’Agostino & Pearson normality test. Paired t-test (two-tailed, assume gaussian distribution), one-way repeated measures ANOVA with Geisser-Greenhouse correction with Sidak’s and Tukey’s correction for multiple comparisons, and two-way repeated measures ANOVA with Sidak’s and Tukey’s correction for multiple comparisons (assume sphericity) was used.

## Data and Code Accessibility

All data and code are freely available through contacting the corresponding author directly.

## ACKNOWLEDGEMENTS

We would like to acknowledge Dr. Pinelopi Kyriazi and Dr. Drew B. Headley from Dr. Denis Pare’s lab for their guidance regarding construction of the active avoidance apparatus and for sharing custom arduino code for running the active avoidance experiments.

## SUPPLEMENTAL FIGURES

**Supplemental Figure 1.**
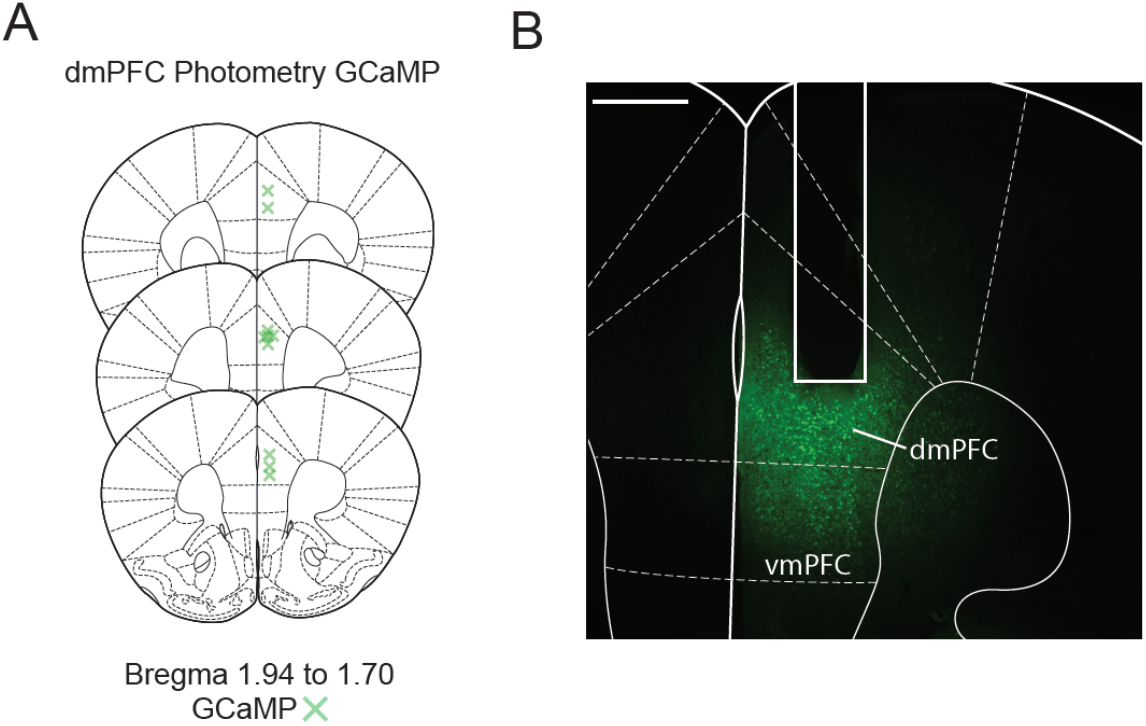
Histology and targeting for dmPFC photometry surgeries. (A) Verification of GCaMP virus injection in dmPFC (N = 10 mice). (B) Representative histological image of fiber photometry implant and GCAMP viral expression in dmPFC. Scale bar 500 μm.

**Supplemental Figure 2.**
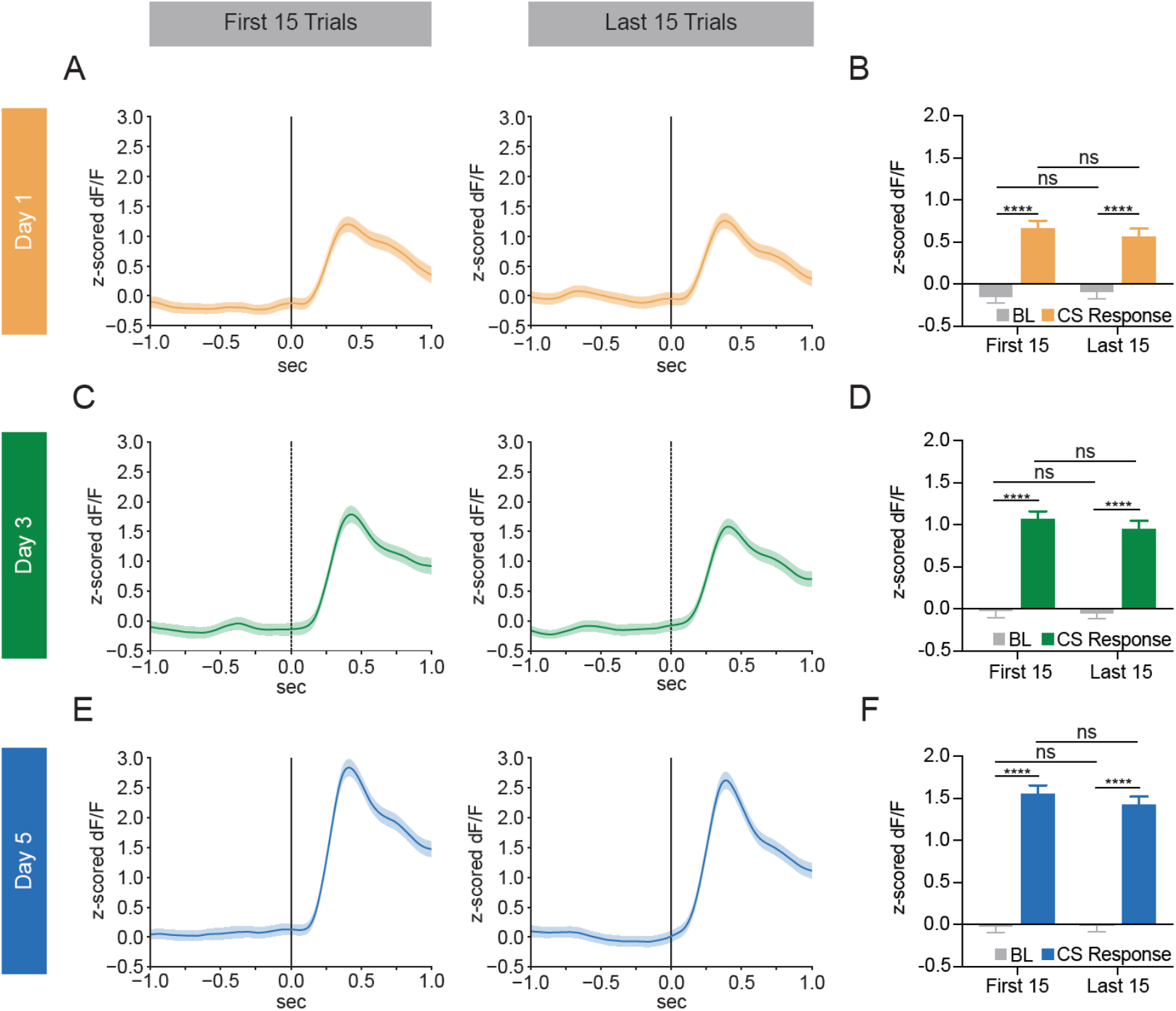
No within-day differences in dmPFC neural activity at CS onset. (A) PETHs of dmPFC calcium signal show no differences between the first 15 trials (left) and the last 15 trials (right) on day 1 of training. (B) Quantification of the day 1 PETHs show no significant differences in calcium signal between the first 15 trials and the last 15 trials during the baseline (−1 to 0 s) or CS (0 to 1 s) periods (Twoway ANOVA, Part of Session x Task Period p = 0.3176, Part of Session p = 0.8385, Task Period p < 0.0001; Sidak’s Multiple Comparisons Test, First 15 Baseline vs Last 15 Baseline p = 0.994, First 15 Baseline vs First 15 CS p < 0.0001, First 15 CS vs Last 15 CS p = 0.9509, Last 15 Baseline vs Last 15 CS p < 0.0001; N = 10 mice, First 15 n = 150 trials, Last 15 n = 150 trials). (C) PETHs of dmPFC calcium signal show no differences between the first 15 trials (left) and the last 15 trials (right) on day 3 of training. (D) Quantification of the day 3 PETHs show no significant differences in calcium signal between the first 15 trials and the last 15 trials during the baseline (−1 to 0 s) or CS (0 to 1 s) periods (Two-way ANOVA, Part of Session x Task Period p = 0.6153, Part of Session p = 0.3854, Task Period p < 0.0001; Sidak’s Multiple Comparisons Test, First 15 Baseline vs Last 15 Baseline p > 0.9999, First 15 Baseline vs First 15 CS p < 0.0001, First 15 CS vs Last 15 CS p = 0.9116, Last 15 Baseline vs Last 15 CS p < 0.0001; N = 10 mice, First 15 n = 150 trials, Last 15 n = 150 trials). (E) PETHs of dmPFC calcium signal show no differences between the first 15 trials (left) and the last 15 trials (right) on day 5 of training. (F) Quantification of the day 5 PETHs show no significant differences in calcium signal between the first 15 trials and the last 15 trials during the baseline (−1 to 0 s) or CS (0 to 1 s) periods (Two-way ANOVA, Part of Session x Task Period p = 0.388, Part of Session p = 0.4610, Task Period p < 0.0001; Sidak’s Multiple Comparisons Test, First 15 Baseline vs Last 15 Baseline p > 0.9999, First 15 Baseline vs First 15 CS p < 0.0001, First 15 CS vs Last 15 CS p = 0.8329, Last 15 Baseline vs Last 15 CS p < 0.0001; N = 10 mice, First 15 n = 150 trials, Last 15 n = 150 trials). ns = not significant, **** p ≤ 0.0001.

**Supplemental Figure 3.**
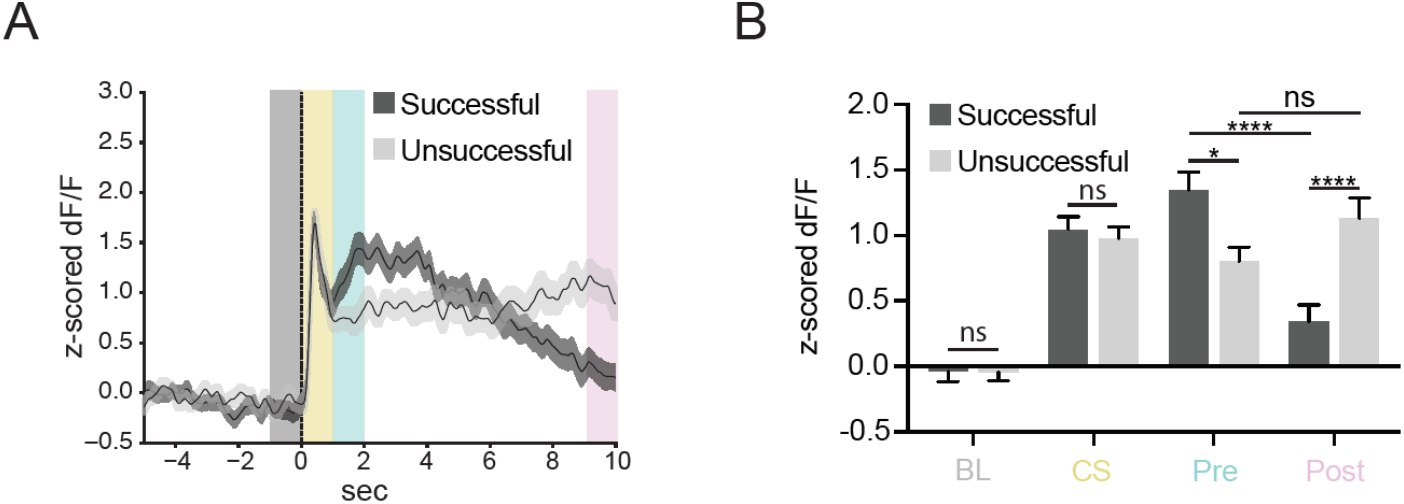
Differences in dmPFC neural activity for successful versus unsuccessful trials. (A) PETH of calcium signal in dmPFC aligned to CS onset for successful (dark grey line) and unsuccessful (light grey line) trials shows differences in later parts of the trace when avoidances normally do or do not occur. Trials from Day 3 were used since equal numbers of successful and unsuccessful trials occur on this training day. Grey box, baseline period (BL); yellow box, CS response period (CS); teal box, pre avoidance period (Pre); pink box, post avoidance period (Post) (B) Quantification of the CS onset PETH shows no differences in calcium signal between successful and unsuccessful trials during the baseline period (−1 to 0 s) and the CS response period (0 to 1 s). However, the calcium signal in the dmPFC is significantly increased during successful trials compared to unsuccessful trials during the pre avoidance period (1 to 2 s) and significantly decreased during successful trials compared to unsuccessful trials during the post avoidance period (9 to 10 s) (Two-way ANOVA, Task Period × Trial Type p < 0.0001, Task Period p < 0.0001, Trial Type p = 0.5807; Sidak’s Multiple Comparisons Test, Successful Baseline vs Unsuccessful Baseline p > 0.9999, Successful CS Response vs Unsuccessful CS Response p = 0.986, Successful Pre Avoidance vs Unsuccessful Pre Avoidance p = 0.0022, Successful Post Avoidance vs Unsuccessful Post Avoidance p < 0.0001; N = 10 mice, Successful n = 147 trials, Unsuccessful n = 153 trials). ns = not significant, * p ≤ 0.05, **** p ≤ 0.0001.

**Supplemental Figure 4.**
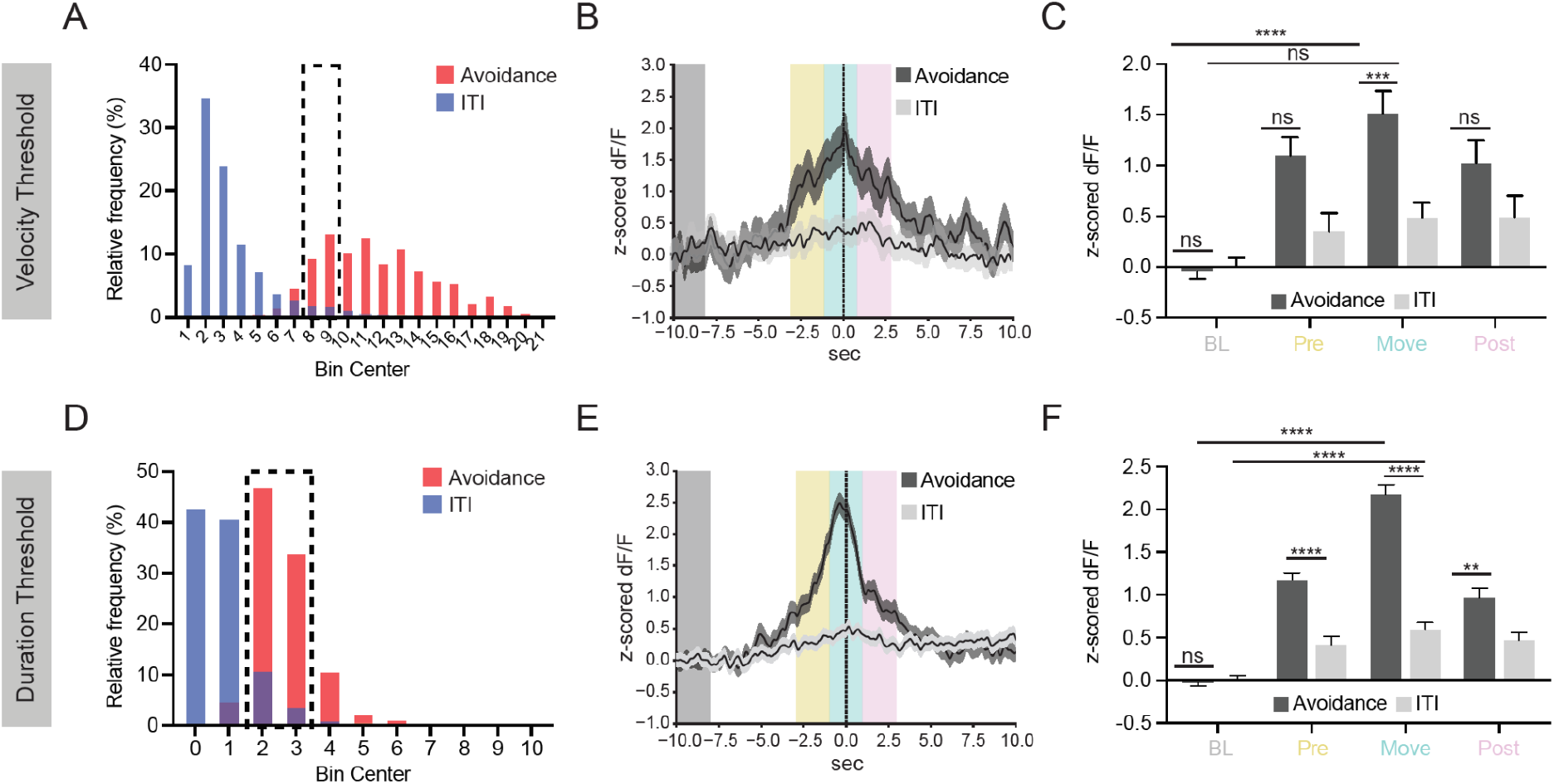
Increased activity in the dmPFC during avoidance is not purely movement-related. (A) Distribution of movement velocities for intertrial (ITI) (blue) and avoidance (red) movements and their overlap (purple). (B) PETH of ITI and avoidance movements of similar velocities (7.5 cm/s to 9.5 cm/s) aligned to movement onset shows increase in calcium signal during avoidance movements that is not seen during ITI movements. Grey box, baseline period (BL); yellow box, pre movement period (Pre); teal box, movement period (Move); pink box, post movement period (Post) (C) Quantification of similar velocity movement PETH show dmPFC calcium signal is significantly increased during avoidance movements compared to ITI movements during the movement period (−1 to 1 s), but not during baseline (−10 to −8 s), pre-movement (−3 to −1 s), or post-movement (1 to 3 s) periods (Two-way ANOVA, Task Period x Movement Type p = 0.015, Task Period p < 0.0001, Movement Type p < 0.0001; Sidak’s Multiple Comparisons Test, Avoidance Baseline vs ITI Baseline p > 0.9999, Avoidance Baseline vs Avoidance Movement p < 0.0001, ITI Baseline vs ITI Movement p = 0.7853, Avoidance Pre-Movement vs ITI Pre-Movement p = 0.0701, Avoidance Movement vs ITI Movement p = 0.0009, Avoidance Post-Movement vs ITI Post-Movement p = 0.5713; N = 10 mice, Avoidance n = 58 trials, ITI n = 60 trials). (D) Distribution of movement durations for ITI (blue) and avoidance (red) movements and their overlap (purple). (E) PETH of ITI and avoidance movements of similar durations (1.5 s to 3.5 s) aligned to movement onset shows sharp increase in calcium signal during avoidance movements that is not seen during ITI movements. (F) Quantification of similar movement duration PETH shows dmPFC calcium signal is significantly increased during avoidance movements compared to ITI movements during pre-movement (−3 to −1 s), movement (−1 to 1 s), and postmovement (1 to 3 s) periods, but not during the baseline (−10 to −8 s) period (Two-way ANOVA, Task Period x Movement Type p < 0.0001, Task Period p < 0.0001, Movement Type p < 0.0001; Sidak’s Multiple Comparisons Test, Avoidance Baseline vs ITI Baseline p > 0.9999, Avoidance Baseline vs Avoidance Movement p < 0.0001, ITI Baseline vs ITI Movement p < 0.0001, Avoidance Pre-Movement vs ITI Pre-Movement p < 0.0001, Avoidance Movement vs ITI Movement p < 0.0001, Avoidance Post-Movement vs ITI Post-Movement p = 0.0019; N = 10 mice, Avoidance n = 205 trials, ITI n = 227 trials). ns = not significant, ** p ≤ 0.01, *** p ≤ 0.001, **** p ≤ 0.0001.

**Supplemental Figure 5.**
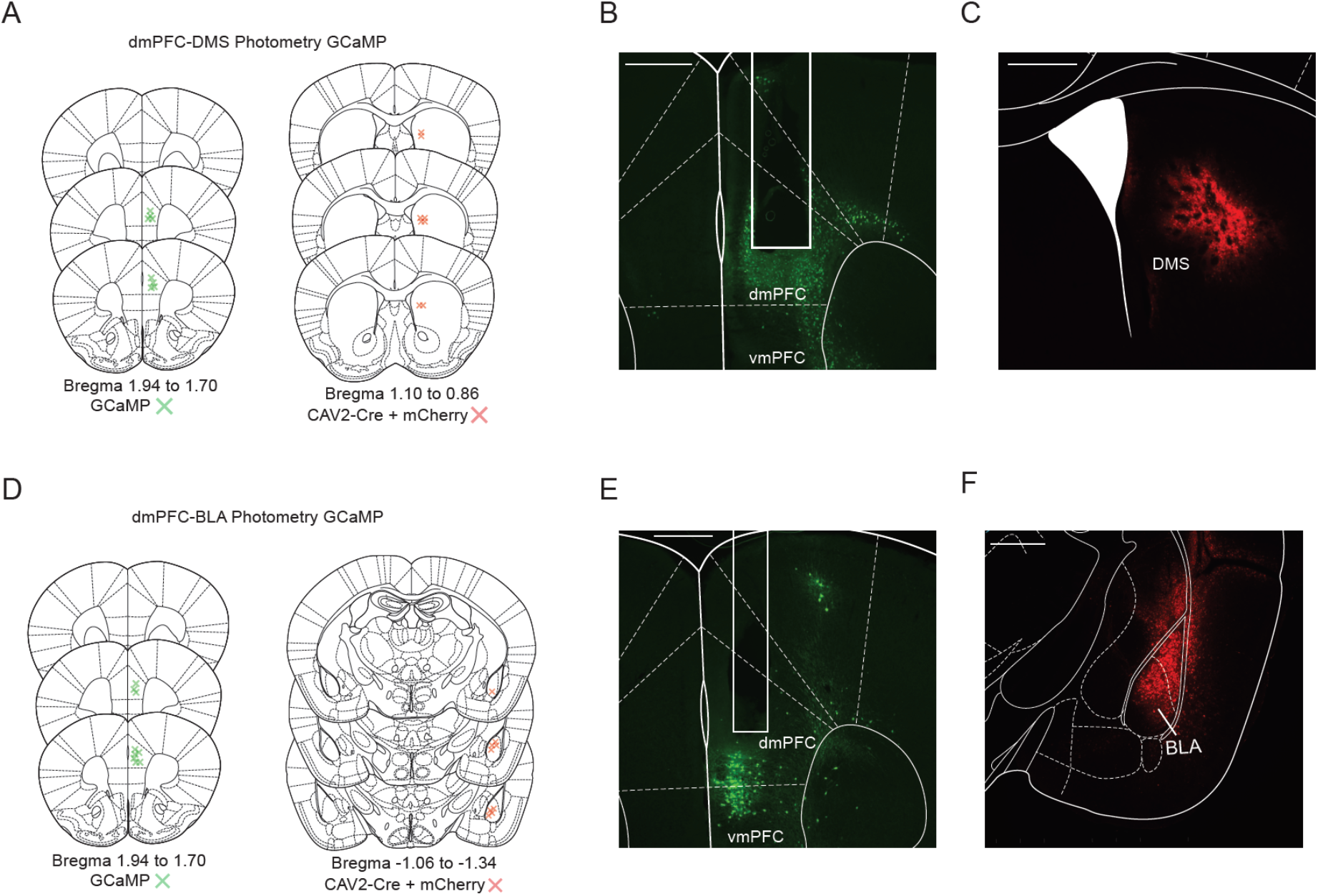
Histology and targeting for dmPFC-DMS and dmPFC-BLA photometry surgeries. (A) Verification of GCaMP virus injection in dmPFC (left) and CAV2-Cre + mCherry viral injection in DMS (right) for dmPFC-DMS cohort (N = 8 mice). (B) Representative histological image of fiber photometry implant and GCAMP viral expression in dmPFC for the dmPFC-DMS cohort. (C) Representative histological image of CAV2-Cre + mCherry viral expression in the DMS for the dmPFC-DMS cohort. (D) Verification of GCaMP virus injection in dmPFC (left) and CAV2-Cre + mCherry viral injection in BLA (right) for dmPFC-BLA cohort (N = 9 mice). (E) Representative histological image of fiber photometry implant and GCAMP viral expression in dmPFC for the dmPFC-BLA cohort. (F) Representative histological image of CAV2-Cre + mCherry viral expression in the BLA for the dmPFC-BLA cohort. Scale bar 500 μm.

**Supplemental Figure 6.**
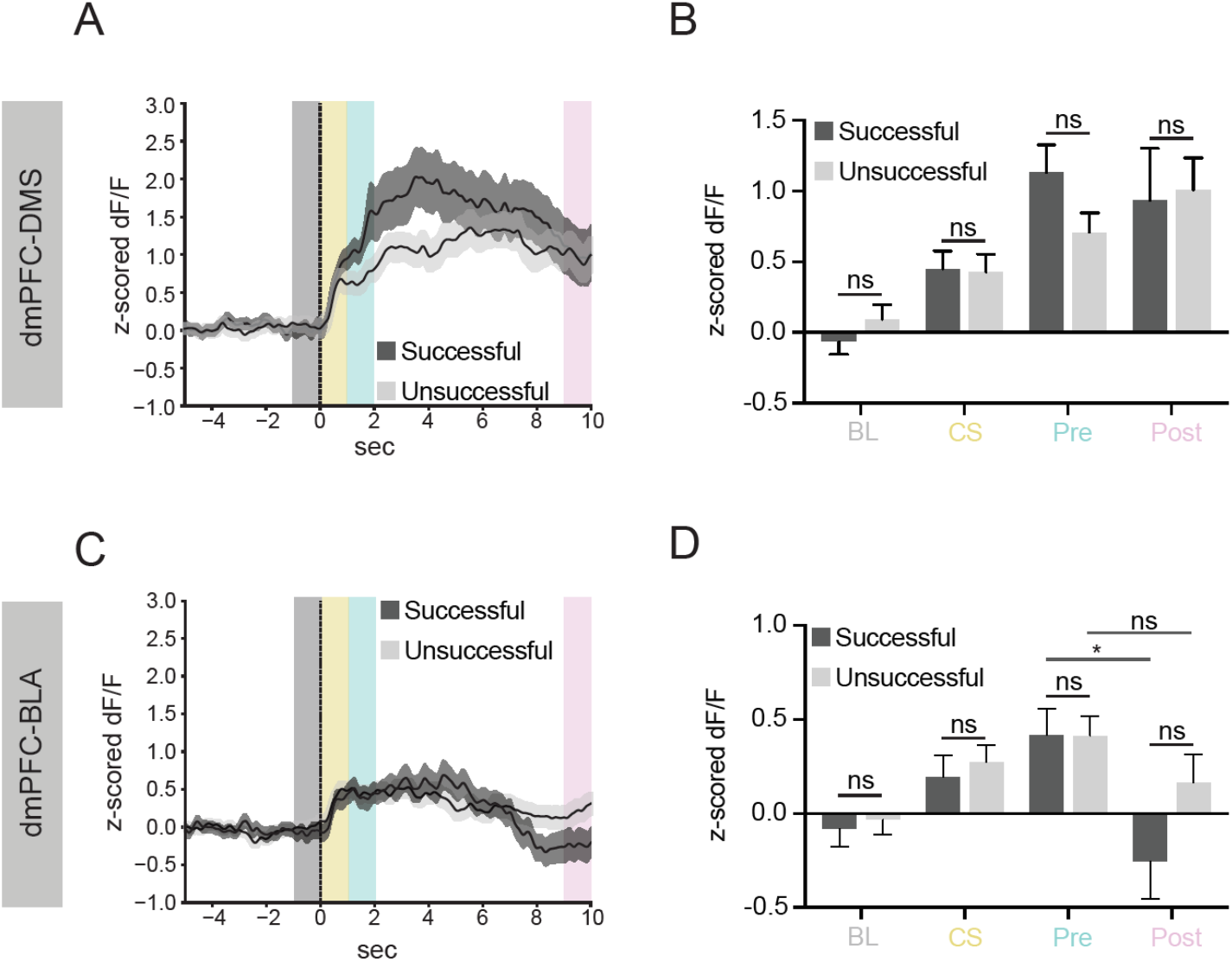
No difference in dmPFC-DMS and dmPFC-BLA neural activity for successful versus unsuccessful trials. (A) PETH of calcium signal in the dmPFC-DMS projection aligned to CS onset for successful (dark grey line) and unsuccessful (light grey line) trials shows no differences between successful and unsuccessful traces. Trials from Day 3 were used since equal numbers of successful and unsuccessful trials occur on this training day. Grey box, baseline period (BL); yellow box, CS response period (CS); teal box, pre avoidance period (Pre); pink box, post avoidance period (Post) (B) Quantification of the CS onset PETH shows no differences in calcium signal between successful and unsuccessful trials during the baseline period (−1 to 0 s), CS response period (0 to 1 s), pre avoidance period (1 to 2 s), or post avoidance period (9 to 10 s) for the dmPFC-DMS projection (Two-way ANOVA, Task Period x Trial Type p < 0.4554, Task Period p < 0.0001, Trial Type p = 0.7025; Sidak’s Multiple Comparisons Test, Successful Baseline vs Unsuccessful Baseline p = 0.9633, Successful CS Response vs Unsuccessful CS Response p > 0.9999, Successful Pre Avoidance vs Unsuccessful Pre Avoidance p = 0.4172, Successful Post Avoidance vs Unsuccessful Post Avoidance p = 0.9978; N = 8 mice, Successful n = 126 trials, Unsuccessful n = 114 trials). (C) PETH of calcium signal in the dmPFC-BLA projection aligned to CS onset for successful (dark grey line) and unsuccessful (light grey line) trials shows no differences between successful and unsuccessful traces on Day 3. (D) Quantification of the CS onset PETH shows no differences in calcium signal between successful and unsuccessful trials during the baseline period (−1 to 0 s), the CS response period (0 to 1 s), the pre avoidance period (1 to 2 s), or the post avoidance period (9 to 10 s) for the dmPFC-BLA projection (Two-way ANOVA, Task Period x Trial Type p = 0.3127, Task Period p < 0.0001, Trial Type p = 0.1204; Sidak’s Multiple Comparisons Test, Successful Baseline vs Unsuccessful Baseline p > 0.9999, Successful CS Response vs Unsuccessful CS Response p > 0.9999, Successful Pre Avoidance vs Unsuccessful Pre Avoidance p > 0.9999, Successful Pre Avoidance vs Successful Post Avoidance p = 0.0137, Unsuccessful Pre Avoidance vs Unsuccessful Post Avoidance p = 0.9679, Successful Post Avoidance vs Unsuccessful Post Avoidance p = 0.3839; N = 9 mice, Successful n = 109 trials, Unsuccessful n = 161 trials). ns = not significant, * p ≤ 0.05.

**Supplemental Figure 7.**
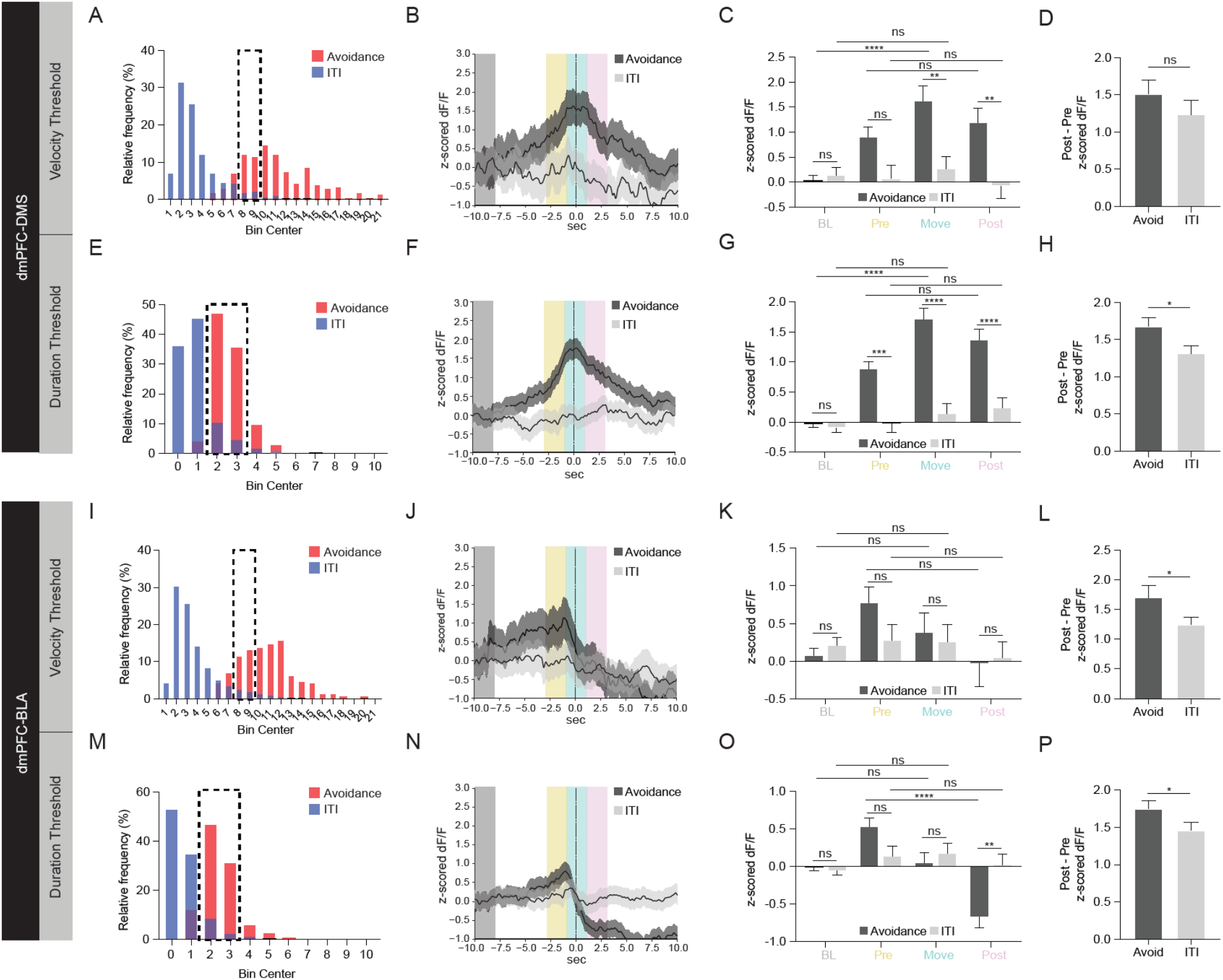
Activity at avoidance onset in the dmPFC-DMS and dmPFC-BLA projections is not purely movement-related. (A) Distribution of movement velocities for ITI (blue) and avoidance (red) movements and their overlap (purple) for the dmPFC-DMS cohort. (B) PETH of ITI and avoidance movements of similar velocities (7.5 cm/s to 9.5 cm/s) aligned to movement onset shows increase in calcium signal during avoidance movements that is not seen during ITI movements in the dmPFC-DMS projection. Grey box, baseline period (BL); yellow box, pre movement period (Pre); teal box, movement period (Move); pink box, post movement period (Post) (C) Quantification of similar velocity movement PETH shows dmPFC-DMS calcium signal is significantly increased during avoidance movements compared to ITI movements during the movement (−1 to 1 s) and post-movement (1 to 3 s) periods, but not during baseline (−10 to −8 s) or pre-movement (−3 to −1 s) periods (Two-way ANOVA, Task Period x Movement Type p 0.0097, Task Period p 0.0049, Movement Type p < 0.0001; Sidak’s Multiple Comparisons Test, Avoidance Baseline vs ITI Baseline p > 0.9999, Avoidance Baseline vs Avoidance Movement p < 0.0001, ITI Baseline vs ITI Movement p > 0.9999, Avoidance Pre-Movement vs ITI Pre-Movement p = 0.3521, Pre-Movement Avoidance vs Post-Movement Avoidance p > 0.9999, Pre-Movement ITI vs Post Movement ITI p > 0.9999, Avoidance Movement vs ITI Movement p = 0.0021, Avoidance Post-Movement vs ITI Post-Movement p = 0.0065; N = 8 mice, Avoidance n = 47 trials, ITI n = 38 trials). (D) To compare the change in the dmPFC-DMS calcium signal between pre-movement and post-movement periods for avoidance and ITI movements, we calculated the absolute value of post-movement change in calcium signal minus pre-movement change calcium signal. We find no differences in the change in dmPFC-DMS calcium signal between pre-movement and post-movement periods when comparing avoidance and ITI movements of similar velocities (Unpaired T-test p = 0.3159; N = 8 mice, Avoidance n = 47 trials, ITI n = 38 trials). (E) Distribution of movement durations for ITI (blue) and avoidance (red) movements and their overlap (purple) for the dmPFC-DMS cohort. (F) PETH of ITI and avoidance movements of similar durations (1.5 s to 3.5 s) aligned to movement onset shows increase in calcium signal during avoidance movements that is not seen during ITI movements in the dmPFC-DMS projection. (G) Quantification of similar movement duration PETH shows dmPFC-DMS calcium signal is significantly increased during avoidance movements compared to ITI movements during the pre-movement (−3 to −1 s), movement (−1 to 1 s), and post-movement (1 to 3 s) periods, but not during the baseline (−10 to −8 s) period (Two-way ANOVA, Task Period x Movement Type p < 0.0001, Task Period p < 0.0001, Movement Type p < 0.0001; Sidak’s Multiple Comparisons Test, Avoidance Baseline vs ITI Baseline p > 0.9999, Avoidance Baseline vs Avoidance Movement p < 0.0001, ITI Baseline vs ITI Movement p = 0.9999, Avoidance Pre-Movement vs ITI Pre-Movement p = 0.0003, Pre-Movement Avoidance vs Post-Movement Avoidance p = 0.3365, Pre-Movement ITI vs Post Movement ITI p = 0.9995, Avoidance Movement vs ITI Movement p < 0.0001, Avoidance Post-Movement vs ITI Post-Movement p < 0.0001; N = 8 mice, Avoidance n = 162 trials, ITI n = 136 trials). (H) The change in the dmPFC-DMS calcium signal between pre-movement and post-movement periods is significantly greater for avoidance movements compared to ITI movements of similar durations (Unpaired T-test p = 0.0166; N = 8 mice, Avoidance n = 162 trials, ITI n = 136 trials). (I) Distribution of movement velocities for ITI (blue) and avoidance (red) movements and their overlap (purple) for the dmPFC-BLA cohort. (J) PETH of ITI and avoidance movements of similar velocities (7.5 cm/s to 9.5 cm/s) aligned to movement onset shows decrease in calcium signal during avoidance movements that is not seen during ITI movements in the dmPFC-BLA projection. (K) Quantification of similar velocity movement PETH shows dmPFC-BLA calcium signal is not significantly different during avoidance movements compared to ITI movements during the baseline (−10 to 8 s), pre-movement (−3 to −1 s), movement (−1 to 1 s), and post-movement (1 to 3 s) periods (Two-way ANOVA, Task Period x Movement Type p = 0.4748, Task Period p = 0.0984, Movement Type p = 0.5066; Sidak’s Multiple Comparisons Test, Avoidance Baseline vs ITI Baseline p > 0.9999, Avoidance Baseline vs Avoidance Movement p > 0.9999, ITI Baseline vs ITI Movement p > 0.9999, Avoidance Pre-Movement vs ITI Pre-Movement p = 0.9625, Pre-Movement Avoidance vs Post-Movement Avoidance p = 0.4489, Pre-Movement ITI vs Post Movement ITI p > 0.9999, Avoidance Movement vs ITI Movement p > 0.9999, Avoidance Post-Movement vs ITI Post-Movement p > 0.9999; N = 9 mice, Avoidance n = 52 trials, ITI n = 88 trials) (L) The change in dmPFC-BLA calcium signal between pre-movement and post-movement periods is significantly greater for avoidance movements compared to ITI movements of similar velocities (Unpaired T-test p = 0.0487; N = 9 mice, Avoidance n = 52 trials, ITI n = 88 trials). (M) Distribution of movement durations for ITI (blue) and avoidance (red) movements and their overlap (purple) for the dmPFC-BLA cohort. (N) PETH of ITI and avoidance movements of similar durations (1.5 s to 3.5 s) aligned to movement onset shows decrease in calcium signal during avoidance movements that is not seen during ITI movements in the dmPFC-BLA projection. (O) Quantification of similar movement duration PETH shows dmPFC-BLA calcium signal is significantly decreased during avoidance movements compared to ITI movements during the post-movement (1 to 3 s) period, but not during the baseline (−10 to −8 s) pre-movement (−3 to −1 s), and movement (−1 to 1 s) periods (Two-way ANOVA, Task Period x Movement Type p = 0.0001, Task Period p < 0.0001, Movement Type p = 0.2688; Sidak’s Multiple Comparisons Test, Avoidance Baseline vs ITI Baseline p > 0.9999, Avoidance Baseline vs Avoidance Movement p > 0.9999, ITI Baseline vs ITI Movement p = 0.9935, Avoidance Pre-Movement vs ITI Pre-Movement p = 0.4972, Pre-Movement Avoidance vs Post-Movement Avoidance p < 0.0001, Pre-Movement ITI vs Post Movement ITI p > 0.9999, Avoidance Movement vs ITI Movement p > 0.9999, Avoidance Post-Movement vs ITI PostMovement p = 0.0017; N = 9 mice, Avoidance n = 165 trials, ITI n = 211 trials). (P) The change in dmPFC-BLA calcium signal between pre-movement and post-movement periods is significantly greater for avoidance movements compared to ITI movements of similar durations (Unpaired T-test p = 0.0473; N = 9 mice, Avoidance n = 165 trials, ITI n = 208 trials). ns = not significant, * p ≥ 0.05, ** p ≤ 0.01, *** p ≤ 0.001, **** p ≤ 0.0001.

